# Sex differences in the regulation and function of cellular immunity in *Drosophila*

**DOI:** 10.1101/2025.03.03.641076

**Authors:** Alexandra Dvoskin, Michael Allara, Kevin Y.L Ho, Nicola Janz, Elizabeth Rideout, Juliet R. Girard, Guy Tanentzapf

**Affiliations:** Department of Cellular and Physiological Sciences, University of British Columbia, Vancouver, V6T 1Z3, Canada; Department of Biology, University of Massachusetts Boston, Boston, Massachusetts, USA; Wyss Institute for Biologically Inspired Engineering, Harvard University, Boston, MA, 02215, USA

## Abstract

Sex differences in development and physiology are prevalent in animals. One physiological system with pronounced differences between the sexes is the immune system: the immune response in humans differs between sexes and results in differential susceptibility of males and females to autoimmune diseases, malignancies, and infectious diseases. However, much remains to be discovered about the mechanisms underlying these sex-based differences in immunity. Here, we use the *Drosophila* hematopoietic organ, the lymph gland, as a model to investigate sex differences in cellular immunity and determine the underlying mechanisms. We find that, in line with their smaller body size, males have smaller lymph glands than females that contain fewer blood progenitors and produce less immune cells. Single cell RNA-seq analysis of the lymph gland showed that they expressed that sex determination genes and identified substantial sex-specific differences in gene expression. By manipulating the sexual identity of different cell types in the lymph gland we show that a subset of these sex differences are controlled by organ-intrinsic mechanisms involving the hematopoietic niche. Importantly, we find a differential response between males and females to changes in insulin signaling, an important regulator of the immune response in the niche. Finally, we provide evidence for differences in the cellular immune response following infection between males and females. Overall, our results provide mechanistic insight into how sex differences in immunity are established.

**Author’s Summary:** Our paper deals with a fundamental question in biology, how does the sex of an organism influence its anatomy and physiology. In particular, we focus on sex-differences in immunity and stem cell function. We establish the Drosophila larva as a model for analysing sex-based differences in cellular immunity. Cellular immunity in Drosophila is based on the production of multiple types of mature immune cells from blood progenitors and takes place in the fly hematopoietic organ, the larval lymph gland. We find that sex controls the number of various cell types in the lymph gland (progenitors, and specific varieties of mature blood cells). These sex-differences vary by cell type and change depending on whether flies are raised under homeostatic or infection conditions. We provide insight into the mechanisms that mediate these sex differences, identifying a possible role for the hematopoietic niche and insulin signaling. Taken together our work not only serves as an initial characterization of baseline sex-differences in fly hematopoiesis and cellular immunity but also identifies important areas for future exploration.

## Introduction

Sex differences exist across many species, where such differences reflect variations in reproductive strategies, ecological roles, and physiological demands [1,2]. In many animals, males and females differ in terms of overall body size, muscle mass, fat storage, and reproductive tissue development [3–13]. These differences often arise early in development, persist throughout life, and manifest at multiple levels, from the individual cell, to tissues, organs, and entire organisms. Sex differences are genetically encoded and typically involve hormonal signals that control animal development and homeostasis [14]. *Drosophila* has proven to be a powerful model for exploring the genetic underpinning and signaling mechanisms that underlie sex differences [15–18]. In *Drosophila*, sex differences arise early, appearing during embryogenesis, and are pronounced in the adults. *Drosophila* males are typically smaller than females, and exhibit marked differences from females in appearance, behavior, and physiology.

It is well established, both in epidemiological and experimental studies, that there are extensive sex differences in immunity in humans[19,20]. This has important implications in health and disease, as evidenced, for example, by sex differences in the ability to fight certain infections, respond to vaccines, or by the higher prevalence of multiple autoimmune conditions in females. It is speculated that sex differences in immunity are attributed to either genetic or hormonal factors and several mechanistic differences in immunity have been uncovered. On a cellular level, innate detection of pathogens is known to differ between males and females, as sex differences have been found in the induction of toll-like receptors (TLR) and the antiviral type I interferon (IFN) response [21,22]. Moreover, in both humans and rats, females show a higher activity of monocytes, macrophages and dendritic cells [23,24]. Females also tend to mount a more robust adaptive immune response than males by having a higher proportion of active T cells during certain viral infections [25], as well as increased expression, associated with estrogen response elements in their promoter regions, of antiviral and inflammatory genes [26].

Although immunology studies in humans have increasingly included sex as a variable, this has not been the case in *Drosophila*, an important genetic model system for immunity. A recent literature survey found that out of over 1000 papers about the adult immune system in *Drosophila,* almost half did not report sex and only 13% reported results for both males and females separately [27]. In the small number of published immunology studies in *Drosophila* that took sex into account, it has been found that there were differences in immune response and survival following infection, with outcomes being pathogen specific. Specifically, viral infections have been found to result in a higher rate of mortality in males than females, with surviving females exhibiting increased infertility [28]. Bacterial infection studies have shown a complex picture, with sex differences in survival rates following infection in certain strains of bacteria, regardless of whether the bacterial strain in question was gram negative or gram positive [29–33]. For example, females were shown to be more susceptible to infection with *Enterococcus faecalis* but less susceptible to *S. aureus* or to *Lactococcus lactis* infection, all of which are gram positive bacteria [27,31,32]. Some of this variation is accounted for by sex-specific differences in the regulation of both IMD and Toll immune pathways between males and females following infection [30,31]. There is also some indirect evidence of sex differences in the cellular immune response, specifically, hemocytes functioning downstream of the Jun N-terminal kinase (JNK) pathway in repairing tissue damage induced by UV irradiation were only seen in males [34,35].

Work in *Drosophila* on hematopoiesis and immunity has focused extensively on the lymph gland, an organ that is responsible for the cellular immune response during larval stages [36]. The primary lobe of the lymph gland is the main site of hematopoiesis in *Drosophila* larval stages and is typically described as being made up of 3 distinct zones: the Posterior Signaling Center (PSC), a group of a few dozen cells that are thought to have a stem cell niche-like function, the medullary zone (MZ), which houses blood progenitors, and the cortical zone (CZ) which holds differentiated blood cells [37].

Multiple signaling pathways act in the lymph gland to regulate hematopoiesis including the Wingless, Hedgehog, JAK/STAT, BMP and Notch pathways [37–42]. Mutations that perturb these signaling pathways result in a variety of defects in lymph gland homeostasis, ranging from depletion of the progenitor population, overproduction of various mature blood cell lineages, and expansion of the PSC [38,39,40,41,43]. Another major signaling pathway known to act in the lymph gland is the insulin/insulin-like growth factor signaling pathway (IIS) [44–46]. In the lymph gland, the IIS pathway has been shown to regulate both the maintenance and differentiation of blood progenitors as well as maintenance of the PSC [47,48]. Moreover, at least two *Drosophila* insulin-like peptides (dILPs), secreted by insulin-producing cells in the brain (dILP-2) and the fat body (dILP-6), are known to act in the lymph gland to control hematopoiesis [49–52].

Here we analyzed sex differences in the *Drosophila* lymph gland under homeostatic conditions as well as following bacterial infection. We identified sex differences in the overall cell number and within specific cell types of the lymph gland. Analysis of single-cell RNA-Seq data from the lymph gland uncovered extensive sex-differences in gene expression in all cell types of the gland. Further investigation into the mechanisms that mediate sex-differences identified a source of some of the relevant regulatory signals, revealing a role for insulin signaling in this process. Finally, we explored whether and how sex differences extend to the cellular immune response following infection.

## Results

### Lymph Glands Exhibit Sex differences

To investigate whether there were sex differences between lymph glands in male and female larvae, we measured organ size in late 3rd instar larva using automated cell counts with custom image analysis software (see Methods). We found that lymph glands from male larvae contained an average of 1587±404 cells (n=59), while lymph glands from females contained on average 2183±501 (n=63), a difference of ∼37% in cell number (Fig. 1F). Next, we determined the number of cells in the Posterior Signaling Centre (PSC), the stem cell niche of the lymph gland by staining with an antibody that labels the transcription factor Antennapedia (Antp) and performing cell counts (see methods). These counts showed that lymph glands from male larvae contained an average of 68±22 PSC cells (n=59), while lymph glands from female larvae contained on average 91±24 PSC cells (n=63), a difference of ∼33% (Fig. 1G).

**Figure 1.**
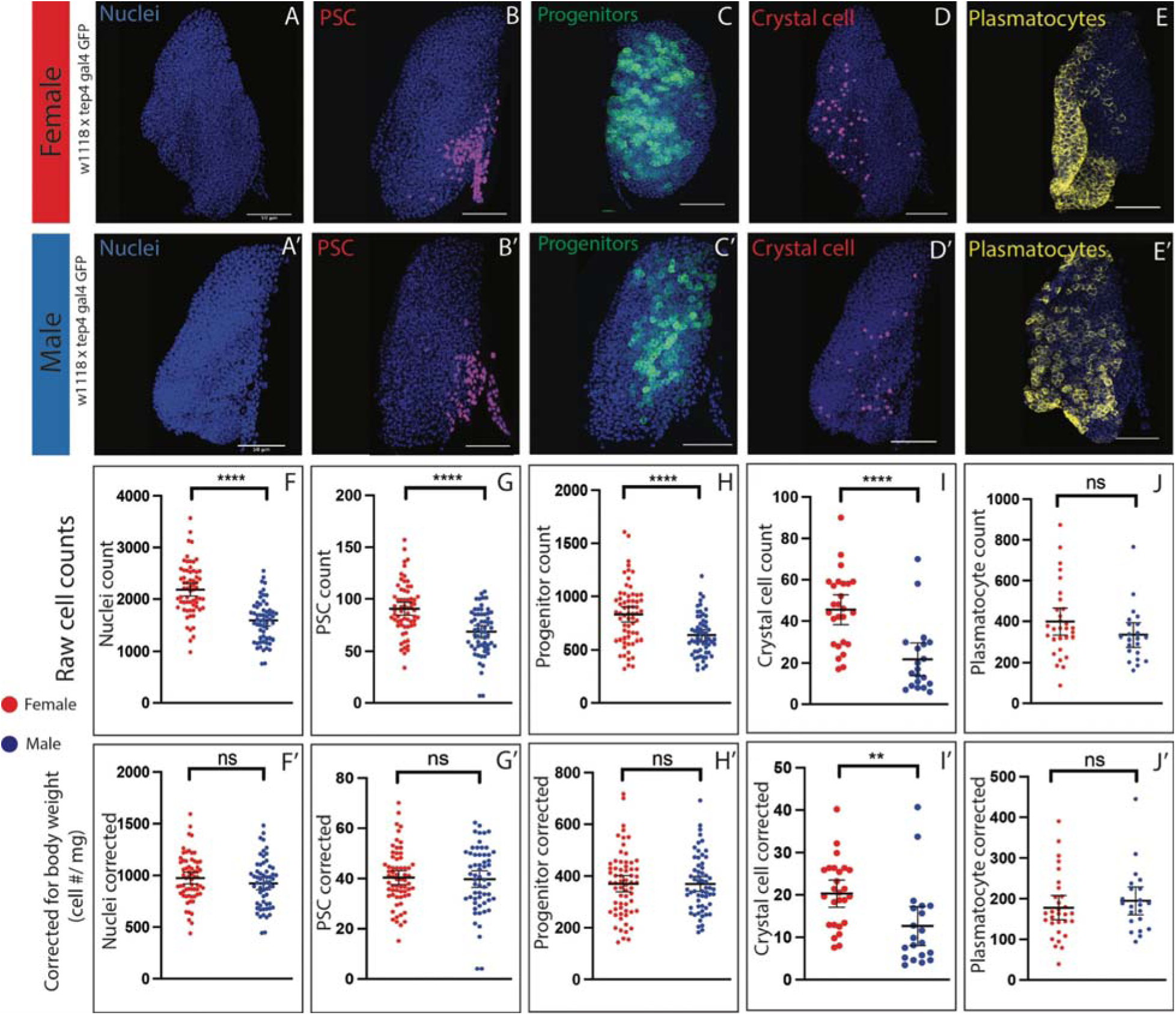
Characterizing sex differences in the lymph gland. (**A-E’**) Representative images of female (A-E) and male (A’-E’) *w^1118^ x tep4-gal4; UAS-GFP* lymph glands stained with ToPro, Antp (B, B’) to mark for PSC cells, endogenously expressing GFP in tep4+ cells (C, C’) to mark progenitors, Hnt (D, D’) to mark for crystal cells, and P1 (E, E’) to mark for plasmatocytes. (F-J) Raw cell counts plotted for both males and females. (F) Raw number of nuclei, used to measure organ size, plotted for males and females (female n=64, male n=59, p<0.0001). (G) Raw number of PSC cells (Antp+), plotted for males and females (female n=64, male n=59, p<0.0001). (H) Raw number of progenitors (tep4+), plotted for males and females (female n=64, male n= 59, p<0.0001). (I) Raw number of crystal cells (Hnt+), plotted for males and females (female n=25, male n=19, p<0.0001). (J) Raw number of plasmatocytes (P1+), plotted for males and females (female n=30, male n=22, p=0.1661). (F’-J’) Corrected for body weight by dividing each female data point by 2.24mg and dividing each male data point by 1.72mg. (F’) Corrected nuclei count, used to measure organ size (female n=64, male n=59, p=0.2107). (G’) Corrected number of PSC cells (Antp+) (female n=64, male n=59, p=0.7819). (H’) Corrected number of progenitors (tep4+) (female n=64, male n=59, p=0,9705). (I’) Corrected number of crystal cells (Hnt+) (female n=25, male n=19, p=0.0056). (J’) Corrected number of plasmatocytes (P1+) (female n=30, male n=22, p=0.4531). **** indicates P<0.0001, *** indicates P<0.001, ** indicates P<0.01, * indicates P<0.05, ns (non-significant) indicates P>0.05. Error bars are 95% CI.

We labelled progenitors in the lymph gland with the marker *domeMESO GFP* to determine the number of cells [51]. Cell counts showed that lymph glands from male larvae contained an average of 636±190 progenitors (n=59), while lymph glands from female larvae contained on average 830±282 progenitors (n=63), a difference of ∼33% (Fig. 1H). To determine the number of mature differentiated plasmatocytes and crystal cells we stained lymph gland with antibodies against the markers P1 and Hindsight (Hnt), respectively, followed by whole lymph gland cell counts (see methods). These cell counts showed that in lymph glands from male larvae there were on average 22±17 crystal cells (n=20) and 335±134 plasmatocytes (n=22), respectively. In comparison, in lymph glands from female larvae there were on average of 46±18 crystal cells (n=28) and 409±172 plasmatocytes (n=29), a difference of ∼210% in crystal cell numbers and ∼22% in plasmatocyte numbers between the sexes (Fig. 1I-J).

Because *Drosophila* male larvae are smaller in size than females, we asked if the sex differences we observed between lymph glands from male and female larvae were consistent with the overall size differences between males and females. To account for known sex differences in body size we normalized lymph gland cell counts to body size (see methods, Fig 1F’-J’). This analysis showed that sex differences in most parameters were eliminated when normalized for body size. In contrast, the female bias in crystal cell number was maintained after normalizing for body size. Taken together, our data shows striking sex differences in both overall organ size as well as the size of specific cell populations in the larval lymph gland. While these differences generally align with expected organ size based on body size, crystal cells show a uniquely large population size increase in females, that exceeds what would be expected based on overall trends in body size.

### RNA sequencing shows differential gene expression between sexes in the lymph gland

The sex-differences in overall organ and cell population size that we observed in the lymph gland raised the possibility of broader, regulatory, differences between male and female lymph glands. To investigate this, we relied on an organ wide single-cell RNA sequencing analysis of lymph glands [53]. The previous work done on this dataset excluded several sex-specific genes prior to graph-based clustering or visualization.

Reanalysis of this data without excluding sex-specific genes (see materials and methods) showed that sex had a significant effect on the UMAP visualization of lymph gland cells. Specifically, we saw physical separation of some of the graph-based cell clusters into two distinct halves within the three-dimensional UMAP (Movie 1). The affected clusters corresponded to lymph gland progenitors (MZ), intermediate cells (intermediate zone or IZ, and proplasmatocyte or proPL), and plasmatocytes (PL; Movie 1). Comparison between the distinct UMAP halves demonstrated that one half is distinguished by high expression of the male-specific long non-coding RNAs (lncRNAs) lncRNA:roX1 and lncRNA:roX2 (Movie 1). Since we added the same number of male and female larvae to each sample, and roughly half of the cells were found to express male-specific genes, this suggested that sex is what distinguishes the two halves of the UMAP.

The differences in lncRNAs were o fparticular interest as it is well established that male flies significantly overexpress roX1 and roX2, which are two long non-coding RNAs involved in X chromosome dosage compensation [54]. roX1 and roX2 are part of the MSL (male specific lethal) complex that binds to and promotes hypertranscription of X-chromosome genes [54]. MSL2 binds to high affinity sites on the X-chromosome which then begins assembly of the full complex including proteins MLE, MSL1, and MOF and roX RNAs [55]. This complex acetylates histone H4 on K16, which promotes additional transcription of genes on the X-chromosome in male flies [55]. One can identify which cells come from male or female flies by differences in these two roX genes. Differences in roX expression are first observed in embryos [56,57] but persist through adulthood and can be seen in multiple larval and adult somatic tissues including the brain [56,58], salivary gland [59], and gut [56].

We disaggregated cells by sex using the expression level of both lncRNA:roX1 and lncRNA:roX2. Cells with high roX1 and roX2 expression (8865 in total, ∼42% of cells) were categorized as male, while cells with low roX1 and roX2 expression (10242 in total, ∼48% of cells) were categorized as female. Cells which showed high expression of one roX RNA but not both (2050 in total, ∼10% of cells) were excluded from further analysis as their sex was ambiguous. When we compared gene expression across the sexes, we observed differential expression of several genes involved in sex determination. For example, we saw that msl-2 was enriched in male cells, but Sxl and tra were enriched in female cells (Figure 2A). These data suggest that the sex determination genes are expressed in lymph gland cells.

**Figure 2.**
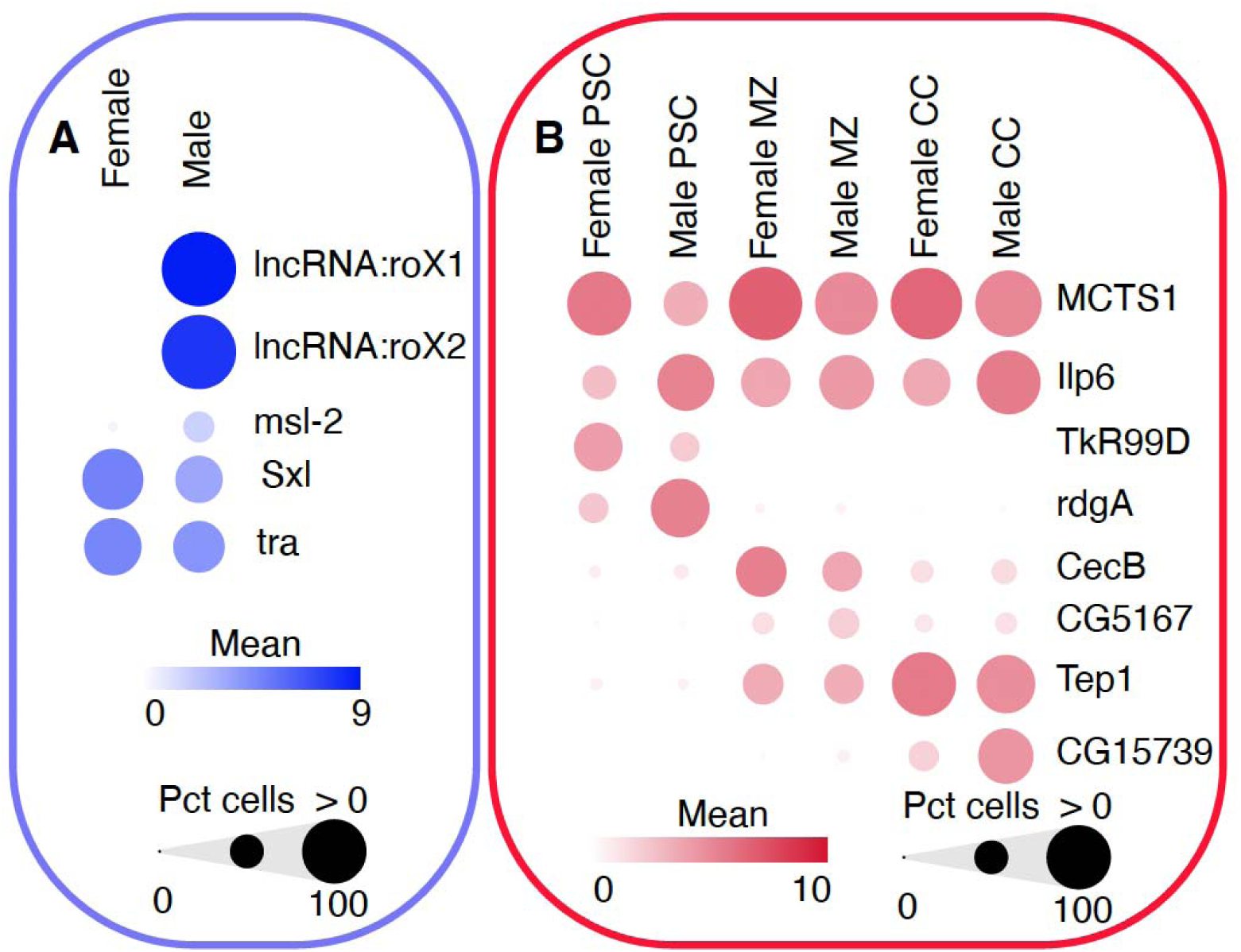
Single-cell RNA sequencing of lymph gland cells. (A-B) Bubble plots showing the mean expression (Mean) of each gene (color intensity) and percentage (Pct) of those cells which express that gene (size). (A) Expression of known sex-specific genes in the scRNA-data that have been disaggregated by sex. lncRNA:roX1, lncRNA:roX2, and msl-2 are enriched in male cells while Sxl and tra are enriched in female cells. (B) Selected genes that were found to be enriched in male or female cells in specific lymph gland zones. PSC, posterior signaling center; MZ, medullary zone; CC, crystal cell.

We identified differentially expressed genes (DEGs) which were significantly enriched in either males or females using ANOVA analysis (genes that were greater than or equal to the fold change threshold of 1.5 and less than or equal to the false discovery rate threshold of 0.0001 were defined as significantly enriched; Supplementary file 1). We then compared gene expression of male and female cells in each graph-based cluster including the crystal cells (CC), PSC, and MZ. This allowed us to determine sex-specific differences within each cell type. While some sex-specific DEGs were differentially expressed across all populations of male or female lymph gland cells, many were only differentially expressed in male or female cells of a specific zone (Supplementary file 1). For example, female PSC cells were uniquely enriched in TkR99D, which encodes a receptor for tachykinin-like neuropeptides, while male PSC cells were specifically enriched for rdgA, a gene encoding a diacylglycerol kinase involved in phospholipase C signaling (Figure 2B). To identify potential pathways or processes involved in each cluster by sex, we performed gene set enrichment analysis on the sex-specific DEGs we identified in each cluster (Supplementary file 1). For example, we found that female MZ cells were enriched in genes involved in the humoral immune response downstream of the Toll and Imd signaling pathways (CecB; Figure 1B; Supplementary file 1). Overall, many of the sex-specific DEGs we identified had human orthologs and some had been previously implicated in blood development, disease, or immune function (Supplementary file 1). These sex-specific genes are of interest for future study and may provide insights into the mechanisms that underlie sex differences in hematopoiesis or immunity.

### Sex differences in the lymph gland are not mediated by the CNS

Sex differences between tissues can arise due to cell-autonomous or non-cell-autonomous mechanisms. For example, it has been shown that systemic signals from tissues such as the fat body, the muscles, or the central nervous system (CNS) can influence cell and body growth in the fly [49–51]. To test whether sex differences in the lymph gland were due to a systemic signal originating from the CNS or fat body, we employed a strategy where we feminized these tissues in an otherwise genetically male fly and asked how this impacted overall organ size as well the size of individual cell populations in the lymph gland. To change the sexual identity of different tissues we expressed sex determination gene *transformer* (*tra*) in males [15,16,60–67]. Normally, a functional Tra protein is only produced in females (Tra^F^), where it specifies most aspects of sexual differentiation and development [68,69,70]. When Tra^F^ is expressed in males, it is sufficient to induce the development of female-specific traits [71]. To control for differences in the lymph gland due to variation in the genetic background, we analysed flies heterozygous for the Tra^F^ construct (Tra^F^/+) and flies heterozygous for the Gal4 driver (Gal4/+) in addition to the experimental group of flies having both the Gal4 and Tra^F^ transgene (Gal4/Tra^F^). We then compared males and females from each of the two control groups as well as the experimental group. This meant we were comparing 6 different genotypes to each other, which required us to use a two-way ANOVA, post-hoc Tukey’s test (see methods) to ask if any differences we observed were statistically significant. This test analyzes the significance of changes between groups and provides a “sex:genotype interaction constant” to determine whether the data supported the existence of sex differences between males and females under the experimental treatment.

When we feminized post-mitotic neurons using *elav*-Gal4, there was no significant effect of genetic background on overall lymph gland size, as males or females of both Tra^F^ and *elav*-Gal4 control flies had a similar number of cells in their lymph gland compared to each other or the experimental group (Figure 3A-D). Moreover, using these controls suggested that CNS feminization had no significant impact on lymph gland size. Indeed, analysing the sex:genotype interaction constant (Table 1) showed that sex differences in the total number of cells in male and female lymph glands were not altered by feminization of the CNS. Similarly, sex differences in the crystal cells or progenitor numbers did not appear to change upon feminization of the CNS, a conclusion that was supported by analysis of the sex:genotype interaction constant (Figure 3E-H and I-L, respectively; Table 1). Taken as a whole, our data and statistical analysis supported the interpretation that sex differences in the lymph gland were not mediated by the CNS.

**Figure 3.**
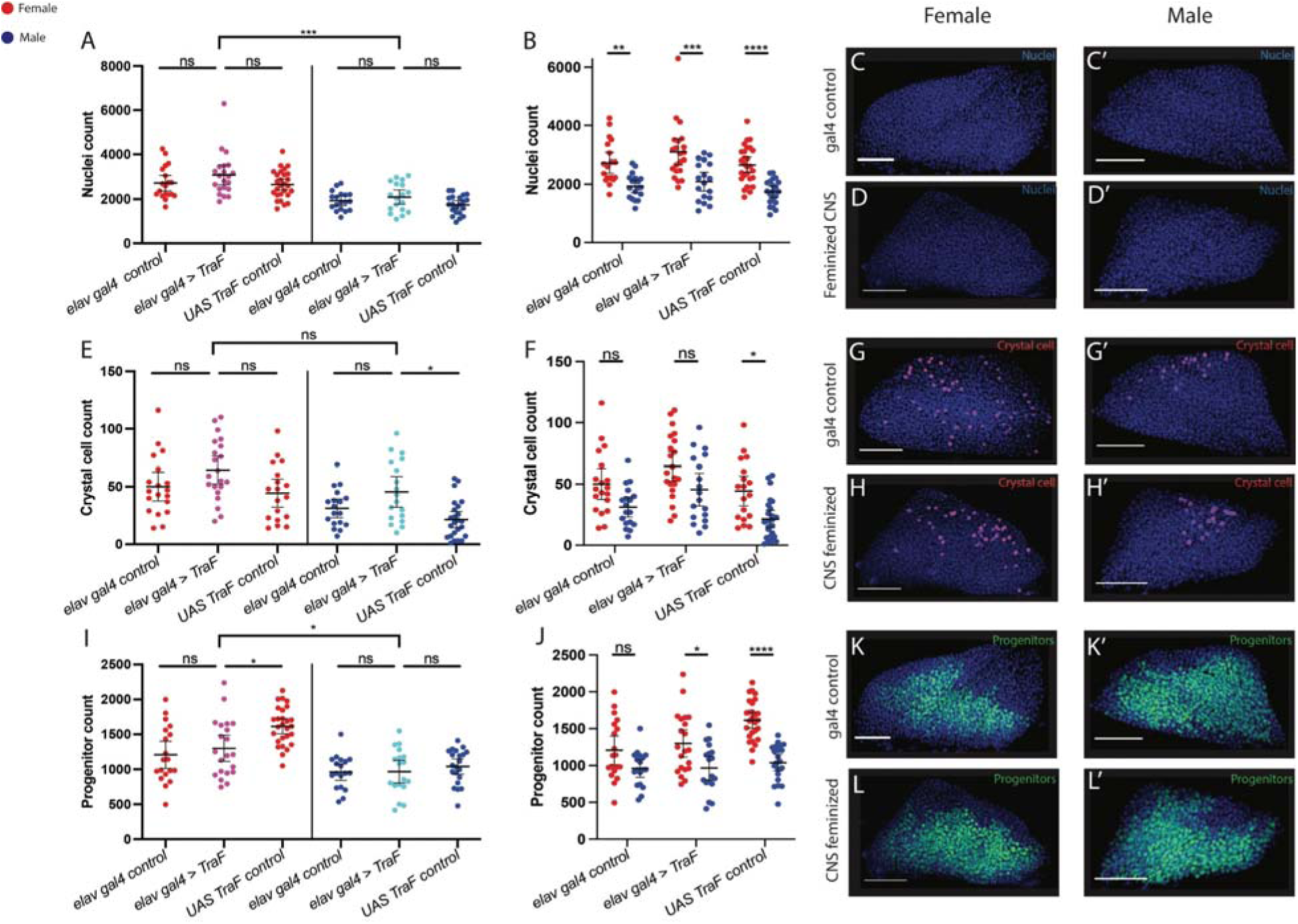
Feminizing the CNS. (A-B) Raw cell counts for total number of nuclei, stained with ToPro. Quantification using a two-way ANOVA, post-hoc Tukey’s test for: *elav-Gal4* control females (n=19), *elav-Gal4>Tra^F^* females (n=21), *UAS Tra^F^* control females (n=27), *elav-Gal4* control males (n=18), *elav-Gal4>Tra^F^* males (n=18), *UAS Tra^F^* control males (n=21). The genotype:sex interaction constant was not significant (p=0.8059). (C) Representative image of *elav-Gal4* control female stained with ToPro. (C’) Representative image of *elav-Gal4* control male stained with ToPro. (D,D’) Representative image of *elav-Gal4;domeMESO GFP > UAS Tra^F^* female (D) and male (D’) stained with ToPro. (E-F) Raw cell counts for total number of crystal cells, stained with Hnt. Quantification using a two-way ANOVA, post-hoc Tukey’s test for: *elav-Gal4* control females(n=19), *elav-Gal4 > Tra^F^*females(n=21), *UAS Tra^F^* control females(n=18), *elav-Gal4* control males(n=18), *elav-Gal4 > Tra^F^* males(n=18), *UAS Tra^F^* control males(n=24) The genotype:sex interaction constant was not significant (p=0.8966). (G, G’) Representative image of *elav-Gal4* control female (G) and male (G’) stained with Hnt. (H, H’) Representative image of *elav-Gal4;domeMESO GFP > UAS Tra^F^*female (H) and male (H’) stained with Hnt. (I-J) Raw cell counts for total number of progenitors, expressing GFP in domeMESO+ cells. Quantification using a two-way ANOVA, post-hoc Tukey’s test, for: *elav-Gal4* control females (n=19), *elav-Gal4 > Tra^F^* females(n=21), *UAS Tra^F^* control females(n=27), *elav-Gal4* control males (n=18), *elav-Gal4 >Tra^F^* males (n=18), *UAS Tra^F^* control males (n=21) The genotype:sex interaction constant was not significant (p=0.0542). (K, K’) Representative image of *elav-Gal4* control female (K) and male (K’), expressing GFP in domeMESO+ cells. (L, L’) Representative image of *elav-Gal4;domeMESO GFP > UAS Tra^F^* female (L) and male (L’), expressing GFP in domeMESO+ cells. **** indicates P<0.0001, *** indicates P<0.001, ** indicates P<0.01, * indicates P<0.05, ns (non-significant) indicates P>0.05. Error bars indicate 95% CI.

**Table 1.** Interaction constants for feminization experiments. Sex:Genotype interaction constants derived from two-way ANOVA, post-hoc Tukey’s test performed on data from Figures 3-6(see methods). Significant interaction constant highlighted in yellow (p<0.05).

|  | Nuclei | Crystal cells | Progenitors |
| --- | --- | --- | --- |
| **Feminized CNS (Fig 3)** | **0.8059** | **0.8966** | **0.0542** |
| **Feminized FB (Fig 4)** | **0.7379** | **0.7910** | **0.0045** |
| **Feminized Progenitors (Fig 5)** | **0.0379** | **0.8740** | **0.0069** |
| **Feminized Niche (Fig 6)** | **0.0139** | **0.0022** | **0.0529** |

### Sex differences in the lymph gland are not mediated by the Fat Body

Given that systemic signals from the fat body can also influence cell and body growth [67,72], we asked whether the sexual identity of this key organ played a role in mediating sex differences in the lymph gland. To this end, we feminized the fat body in genetically male flies using the Tra^F^ transgene and studied how this impacted individual cell population or overall organ size in the lymph gland. As we did for the CNS feminization experiments, we compared experimental groups of flies having both the *R4*-Gal4 and Tra^F^ transgene (*R4*-Gal4/Tra^F^) to control flies heterozygous for either the Tra^F^ construct (Tra^F^/+) or the Gal4 driver (*R4*-Gal4/+). We then performed statistical analysis by calculating the sex:genotype interaction constant (Table 1) to determine if any sex differences in the total number of cells, crystal cells, or progenitors in male and female lymph glands were altered by feminization of the fat body in males.

Our control experiments showed that in the genetic background of *R4*-Gal4, the driver we employed for expression in the fat body, the overall size of the lymph gland was smaller than other backgrounds (Figure 4A). However, since this effect was seen in both males and females it did not impact our interpretation. In particular, we found that feminization of the fat body had no significant impact on size differences between male and female lymph glands (Figure 4A-D, table 1). In contrast to what was observed for overall organ size, crystal cell numbers were higher in lymph glands of the experimental group compared to Tra^F^ and *R4*-Gal4 controls for both males and females.

**Figure 4.**
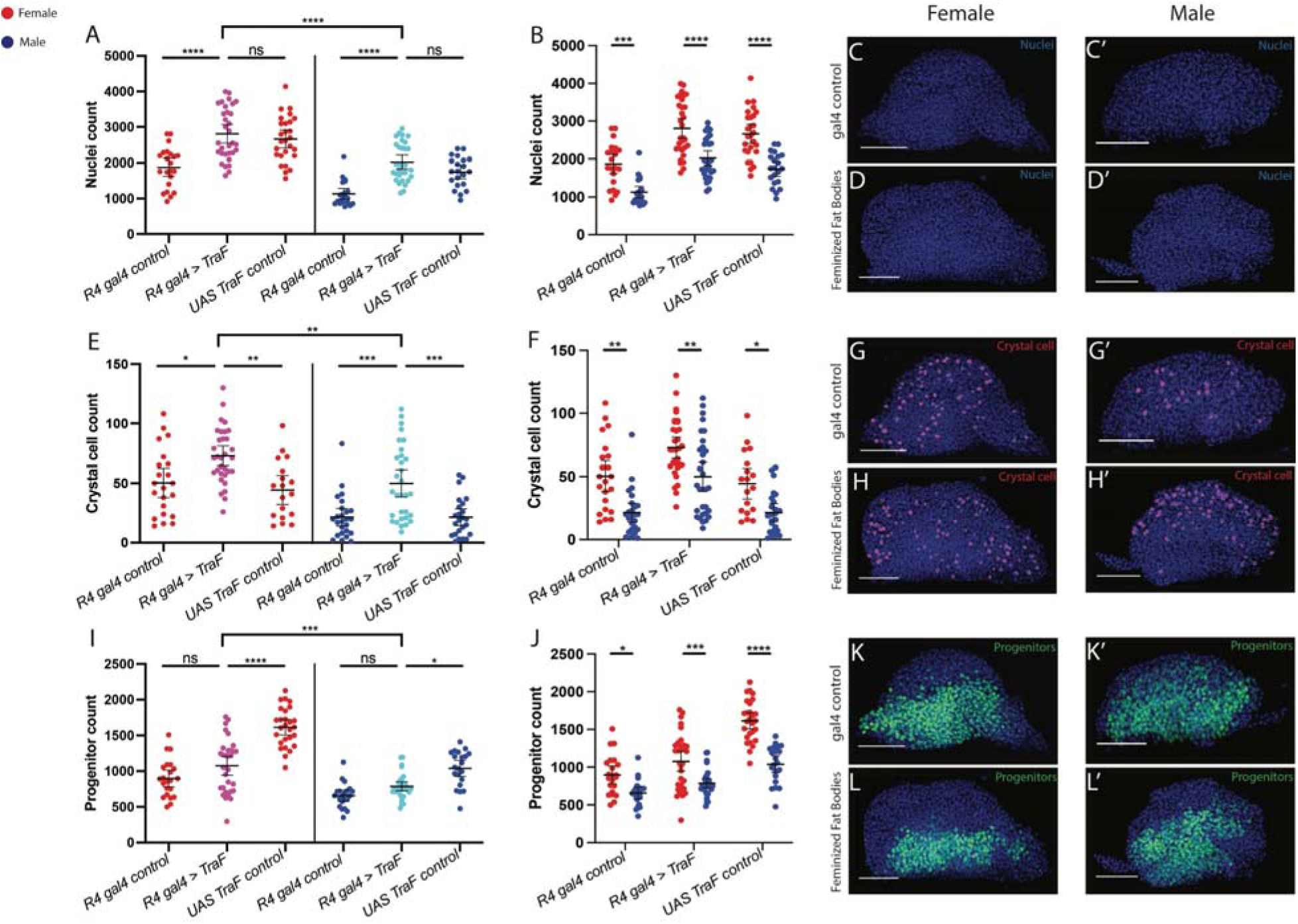
Feminizing the fat bodies. (A-B) Raw cell counts for total number of nuclei, stained with ToPro. Quantification using a two-way ANOVA, post-hoc Tukey’s test, for: *R4-Gal4* female control (n=22), *R4-Gal4* feminized females (n=32), *UAS Tra^F^*control females (n=27), *R4-Gal4* control males (n=21),*R4-Gal4* feminized males (n=31), *UAS Tra^F^* control males (n=21). The genotype:sex interaction constant was not significant (p=0.7379). (C, C’) Representative image of *R4-Gal4* control female (C) and male (C’) stained with Topro. (D) Representative image of *R4-Gal4;domeMESO GFP < UAS Tra^F^*feminized female stained with ToPro. (D’) Representative image of *R4-Gal4;domeMESO GFP < UAS Tra^F^* feminized male stained with ToPro. (E-F) Raw cell counts for total number of crystal cells, stained with Hnt. Quantification using a two-way ANOVA, post-hoc Tukey’s test, for: *R4-Gal4* control (n=22), *R4-Gal4* feminized females (n=32), *UAS Tra^F^* control (n=18), *R4-Gal4* control males (n=25), *R4-Gal4* feminized males (n=31), *UAS Tra^F^* control males (n=24). The genotype:sex interaction constant was no significant (p=0.7910). (G, G’) Representative image of *R4-Gal4* control female (G) and male (G’) stained with Hnt. (H) Representative image of *R4-Gal4;domeMESO GFP x UAS Tra^F^* feminized female stained with Hnt. (H’) Representative image of *R4-Gal4;domeMESO GFP x UAS Tra^F^* feminized male stained with Hnt. (I-J) Raw cell counts for total number of progenitors, expressing GFP in domeMESO+ cells. Quantification using a two-way ANOVA, post-hoc Tukey’s test, for: *R4-Gal4* control females (n=22), *R4-Gal4* feminized females (n=32), *UAS Tra^F^*control females(n=27), *R4-Gal4* control males (n=21), *R4-Gal4* feminized males (n=31), *UAS Tra^F^* control males(n=21). The genotype:sex interaction constant was significant (p=0.0045). (K, K’) Representative image of *R4-Gal4* control female (K) and male (K’), expressing GFP in domeMESO+ cells. (L) Representative image of *R4-Gal4;domeMESO GFP x UAS Tra^F^* feminized female, expressing GFP in domeMESO+ cells. (L’) Representative image of *R4-Gal4;domeMESO GFP x UAS Tra^F^* feminized male, expressing GFP in domeMESO+ cells.**** indicates P<0.0001, *** indicates P<0.001, ** indicates P<0.01, * indicates P<0.05, ns (non-significant) indicates P>0.05. Error bars indicate 95% CI.

Nonetheless, despite this increase, we found no significant impact on sex differences in crystal cell numbers upon fat body feminization, as the increase was observed in both males and females (Figure 4E-H, Table 1). When progenitor numbers were analysed, we observed a statistically significant increase in their number in the background of the Tra^F^ transgene (Figure 3I). Nonetheless, there was no clear impact on sex differences in progenitor numbers (Figure 4I-L, Table 1). Taken as a whole, our data and statistical analysis supported the interpretation that sex differences in overall size, progenitor number, and crystal cell numbers were not mediated by the fat body.

### Sex differences in the lymph gland are not mediated by progenitors

Since we did not uncover compelling evidence for a role of systemic signals from the CNS or the fat body in mediating sex differences in the lymph gland we shifted our focus to lymph gland specific mechanisms. We asked whether the identity of the blood progenitors themselves controlled sex differences. To test this hypothesis, we feminized the blood progenitors in genetically male flies using the Tra^F^ transgene and studied how this impacted overall size as well as the size of individual cell populations in the lymph gland. As before, we compared experimental group flies to flies heterozygous for either the Tra^F^ construct (Tra^F^/+) or the Gal4 driver (*tep4*-Gal4/+) and performed statistical analysis by determining the sex:genotype interaction constant (Table 1).

We observed no effect from changing the sexual identity of blood progenitors in genetically male flies on lymph gland size or on crystal cell number (Fig. 5A-D and E-H, respectively; Table1). While there was a decrease in progenitor cell counts in females with progenitor-specific Tra^F^ expression, there was no significant impact on progenitor numbers in feminized males (Fig 5I-L, Table 1). Taken as a whole, our data shows that feminizing lymph gland progenitors does not allow males to achieve a larger lymph gland, a higher number of crystal cells, or an increase in progenitor cell numbers. Overall, these experiments did not support a significant role for progenitors in mediating sex differences in the lymph gland.

**Figure 5.**
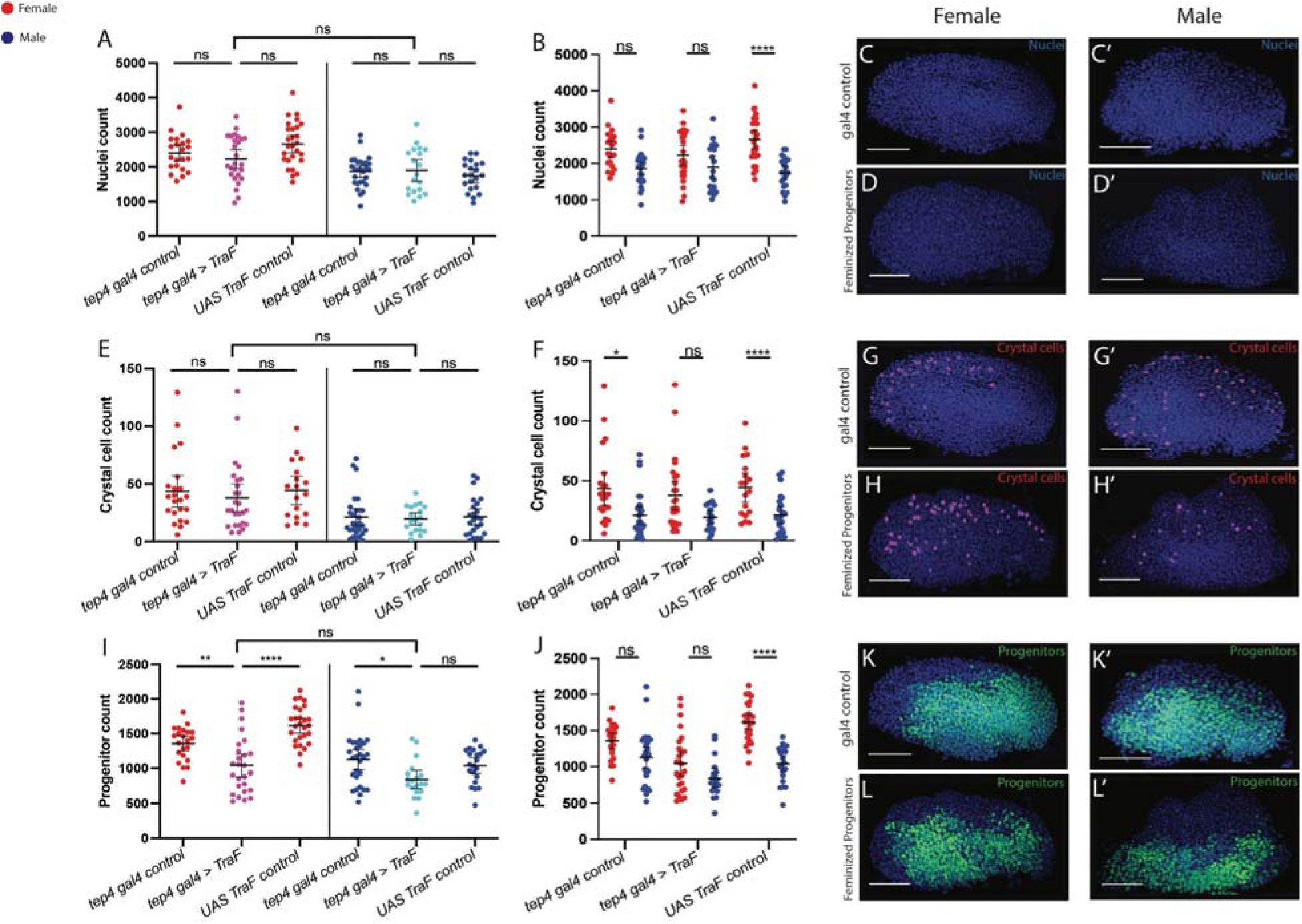
Feminizing the progenitors. (A-B) Raw cell counts for total number of nuclei, stained with ToPro. Quantification using a two-way ANOVA, post-hoc Tukey’s test, for: *tep4-Gal4* control females (n=22), *tep4-Gal4* feminized females (n=26), *UAS Tra^F^*control females (n=27), *tep4-Gal4* control males (n=28), *tep4-Gal4* feminized males (n=19), *UAS Tra^F^* control males (n=21). The genotype:sex interaction constant was significant (p=0.0379). (C, C’) Representative image of *tep4-Gal4* control female (C) and male (C’) stained with Topro. (D, D’) Representative image of *tep4-Gal4;domeMESO GFP < UAS Tra^F^* feminized female (D) and male (D’) stained with ToPro. (E-F) Raw cell counts for total number of crystal cells, stained with Hnt. Quantification using a two-way ANOVA, post-hoc Tukey’s test, for: *tep4-Gal4* control females (n=22), *tep4-Gal4* feminized females (n=26), *UAS Tra^F^* control females (n=18), *tep4-Gal4* control males (n=28), *tep4-Gal4* feminized males (n=19), *UAS Tra^F^* control males (n=24). The genotype:sex interaction constant was not significant (p=0.8740). (G) Representative image of *tep4-Gal4* control female stained with Hnt. (G’) Representative image of *tep4gal4* control male stained with Hnt. (H) Representative image of *tep4-Gal4;domeMESO GFP x UAS Tra^F^* feminized female stained with Hnt. (H’) Representative image of *tep4-Gal4;domeMESO GFP x UAS Tra^F^* feminized male stained with Hnt. (I-J) Raw cell counts for total number of progenitors, expressing GFP in domeMESO+ cells. Quantification using a two-way ANOVA, post-hoc Tukey’s test, for: *tep4-Gal4* control females (n=22), *tep4-Gal4* feminized females (n=26), *UAS Tra^F^* control females (n=27), t*ep4-Gal4* control males (n=28), *tep4-Gal4* feminized males (n=19), *UAS Tra^F^*control males (n=21). The genotype:sex interaction constant was significant (p=0.0069) (K) Representative image of *tep4-Gal4* control female, expressing GFP in domeMESO+ cells. (K’) Representative image of *tep4-Gal4* control male, expressing GFP in domeMESO+ cells. (L) Representative image of *tep4-Gal4;domeMESO GFP x UAS Tra^F^* feminized female, expressing GFP in domeMESO+ cells. (L’) Representative image of *tep4-Gal4;domeMESO GFP x UAS Tra^F^* feminized male, expressing GFP in domeMESO+ cells. **** indicates P<0.0001, *** indicates P<0.001, ** indicates P<0.01, * indicates P<0.05, ns (non-significant) indicates P>0.05. Error bars indicate 95% CI.

### Some sex differences in the lymph gland are mediated by the PSC

The PSC has an established role in orchestrating the behaviour of different cell types in the lymph gland and is therefore a candidate for mediating sex differences. Intriguingly, feminization of the PSC in males, using *collier*-Gal4, induced an increase in the number of cells in the lymph gland which led, for the most part, to the elimination of sex differences in size between females and PSC-feminized males (Figure 6A-D, Table 1). A similar effect was seen for crystal cell numbers, which were higher in PSC-feminized males and resembled control females (Figure 6E-H, Table 1). Thus, the sex difference in both overall lymph gland size and crystal cell number was eliminated due to a male-specific increase in these parameters with Tra^F^ expression (sex:genotype interaction p= 0.0139 and 0.0022, respectively; two-way ANOVA with post-hoc Tukey’s test).

**Figure 6.**
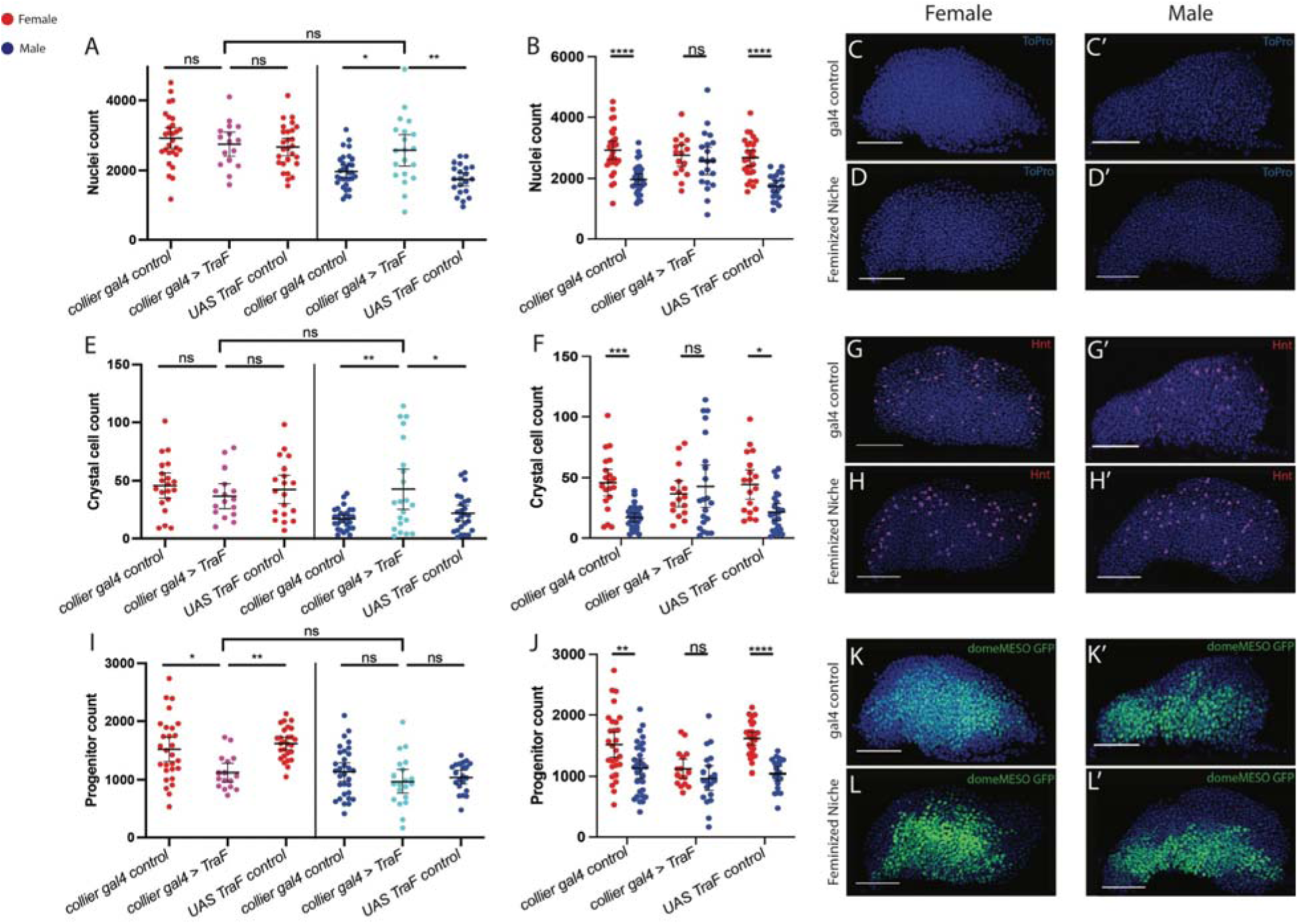
Feminizing the PSC. (A-B) Raw cell counts for total number of nuclei, stained with ToPro. Quantification using a two-way ANOVA, post-hoc Tukey’s test, for: *collier-Gal4* control females (n=28), *collier-Gal4* feminized females (n=16), *UAS Tra^F^* control females (n=27), *collier-Gal4* control males(n=31), *collier-Gal4* feminized males(n=20), *UAS Tra^F^* control males (n=21). The genotype:sex interaction constant was significant (p=0.0139). (C, C’) Representative image of *collier-Gal4* control female (C) and male (C’) stained with Topro. (D) Representative image of *collier-Gal4;domeMESO GFP < UAS Tra^F^* feminized female stained with ToPro. (D’) Representative image of *collier-Gal4;domeMESO GFP < UAS Tra^F^* feminized male stained with ToPro. (E-F) Raw cell counts for total number of crystal cells, stained with Hnt. Quantification using a two-way ANOVA, post-hoc Tukey’s test, for: *collier-Gal4* control females (n=20), *collier-Gal4* feminized females (n=16), *UAS Tra^F^* control females (n=18), *collier-Gal4* control males (n=29), *collier-Gal4* feminized males (n=21), *UAS Tra^F^* control males (n=24). The genotype:sex interaction constant was significant (p=0.0022). (G, G’) Representative image of *collier-Gal4* control female (G) and male (G’) stained with Hnt. (H) Representative image of *collier-Gal4;domeMESO GFP x UAS Tra^F^* feminized female stained with Hnt. (H’) Representative image of *collier-Gal4;domeMESO GFP x UAS Tra^F^* feminized male stained with Hnt. (I-J) Raw cell counts for total number of progenitors, expressing GFP in domeMESO+ cells. Quantification using a two-way ANOVA, post-hoc Tukey’s test, for: *collier-Gal4* control females (n=28), *collier-Gal4* feminized females (n=16), *UAS Tra^F^* control females (n=27), *collier-Gal4* control males (n=31), *collier-Gal4* feminized males (n=20), *UAS Tra^F^* control males (n=21) and *collier-Gal4* feminized males (p=0.9910). The genotype:sex interaction constant was not significant (p=0.0529). (K, K’) Representative image of *collier-Gal4* control female (K) and male (K’), expressing GFP in domeMESO+ cells. (L) Representative image of *collier-Gal4;domeMESO GFP x UAS Tra^F^* feminized female, expressing GFP in domeMESO+ cells. (L’) Representative image of *collier-Gal4;domeMESO GFP x UAS Tra^F^* feminized male, expressing GFP in domeMESO+ cells.**** indicates P<0.0001, *** indicates P<0.001, ** indicates P<0.01, * indicates P<0.05, ns (non-significant) indicates P>0.05. Error bar indicates 95% CI.

In contrast to overall size or crystal cell numbers, there was a significant decrease in progenitor cell counts in females with no effect in males (Figure 6I-L). This effect was similar between the sexes as the sex:genotype interaction was not significant (p= 0.0529; two-way ANOVA). Nonetheless, our data and statistical analysis did support the interpretation that sex differences in overall lymph gland size and crystal cell numbers were mediated, at least in part, by the PSC.

### Sex differences in crystal cell number are mediated by insulin signaling in the PSC

Insulin signaling is known to be an important mediator of sex differences in size and tissue growth [73–76]. Insulin signaling is also known to be active in the PSC in the lymph gland to regulate hematopoiesis [50,51]. Based on these observations we hypothesized that insulin signaling, in the PSC, was important in mediating sex differences in the lymph gland. For these experiments we used *collier*-Gal4 to express either an RNAi to knock down the insulin receptor (InR-RNAi, Figure 7) or a constitutively active version of the insulin receptor (Ca-InR, Figure 8).

**Figure 7.**
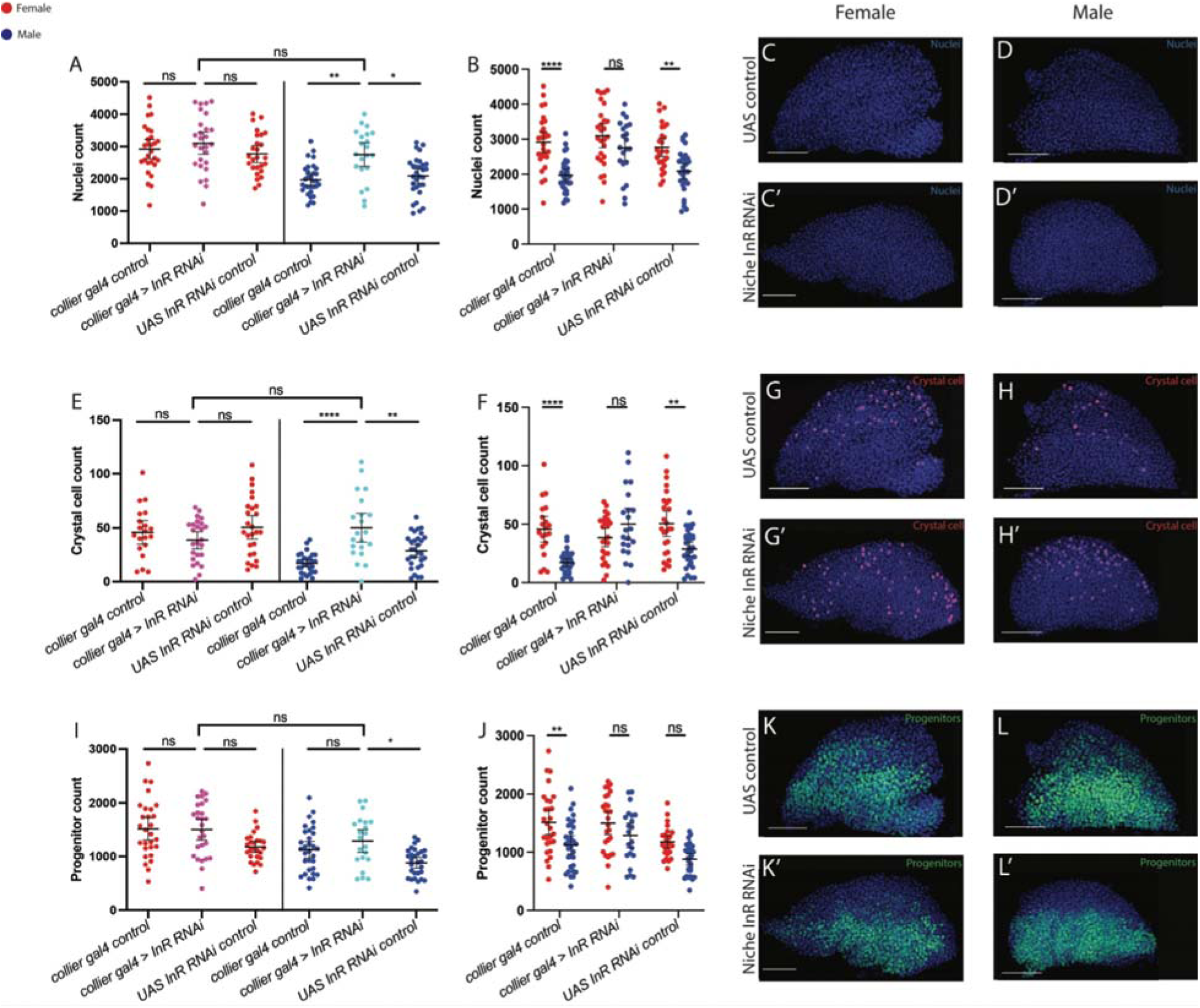
Knocking down insulin signaling in the PSC. (A-B) Raw cell counts for total number of nuclei, stained with ToPro. Quantification using a two-way ANOVA, post-hoc Tukey’s test, for: *collier-Gal4* control females (n=28), *collier-Gal4>InR RNAi* females (n=27), *UAS InR RNAi* females (n=26), *collier-Gal4* control males (n=31), *collier-Gal4>InR RNAi* males (n=21), *UAS InR RNAi* control males (n=30). The sex:genotype interaction constant was not significant (p=0.0931). (C) Representative image of *+> UAS InR RNAi* control females stained with ToPro. (C’) Representative image of *collier-Gal4 > UAS InR RNAi* female stained with ToPro. (D) Representative image of *+> UAS InR RNAi* control male stained with ToPro, (D’) Representative image of *collier-Gal4 > UAS InR RNAi* male stained with ToPro. (E-F) Raw cell counts for total number of crystal cells, stained with Hnt. A two-way ANOVA, post-hoc Tukey’s test, for: *collier-Gal4* control females (n=20), *collier-Gal4>InR RNAi* females (n=25), *UAS InR RNAi* females (n=26), *collier-Gal4* control males (n=30), *collier-Gal4>InR RNAi* males (n=21), *UAS InR RNAi* control males (n=30). The sex:genotype interaction constant was significant (p<0.0001). (G) Representative image of *+> UAS InR RNAi* control females stained with Hnt. (G’) Representative image of *collier-Gal4 > UAS InR RNAi* female stained with Hnt. (H) Representative image of *+> UAS InR RNAi* control male stained with Hnt (H’) Representative image of *collier-Gal4 > UAS InR RNAi* male stained with Hnt. (I-J) Raw cell counts for total number of progenitors, expressing GFP in domeMESO+ cells. A two-way ANOVA, post-hoc Tukey’s test, for: *collier-Gal4* control females (n=28), *collier-Gal4>InR RNAi* females (n=27), *UAS InR RNAi* females (n=26), *collier-Gal4* control males (n=31), *collier-Gal4>InR RNAi* males (n=21), *UAS InR RNAi* control males (n=30). The genotype:sex interaction constant was not significant (p=0.5419). (K) Representative image of *+> UAS InR RNAi* control females expressing GFP in domeMESO+ cells. (K’) Representative image of *collier-Gal4 > UAS InR RNAi* female expressing GFP in domeMESO+ cells. (L) Representative image of *+> UAS InR RNAi* control male expressing GFP in domeMESO+ cells. (L’) Representative image of *collier gal4 > UAS InR RNAi* male expressing GFP in domeMESO+ cells.**** indicates P<0.0001, *** indicates P<0.001, ** indicates P<0.01, * indicates P<0.05, ns (non-significant) indicates P>0.05. Error bar indicates 95% CI.

**Figure 8.**
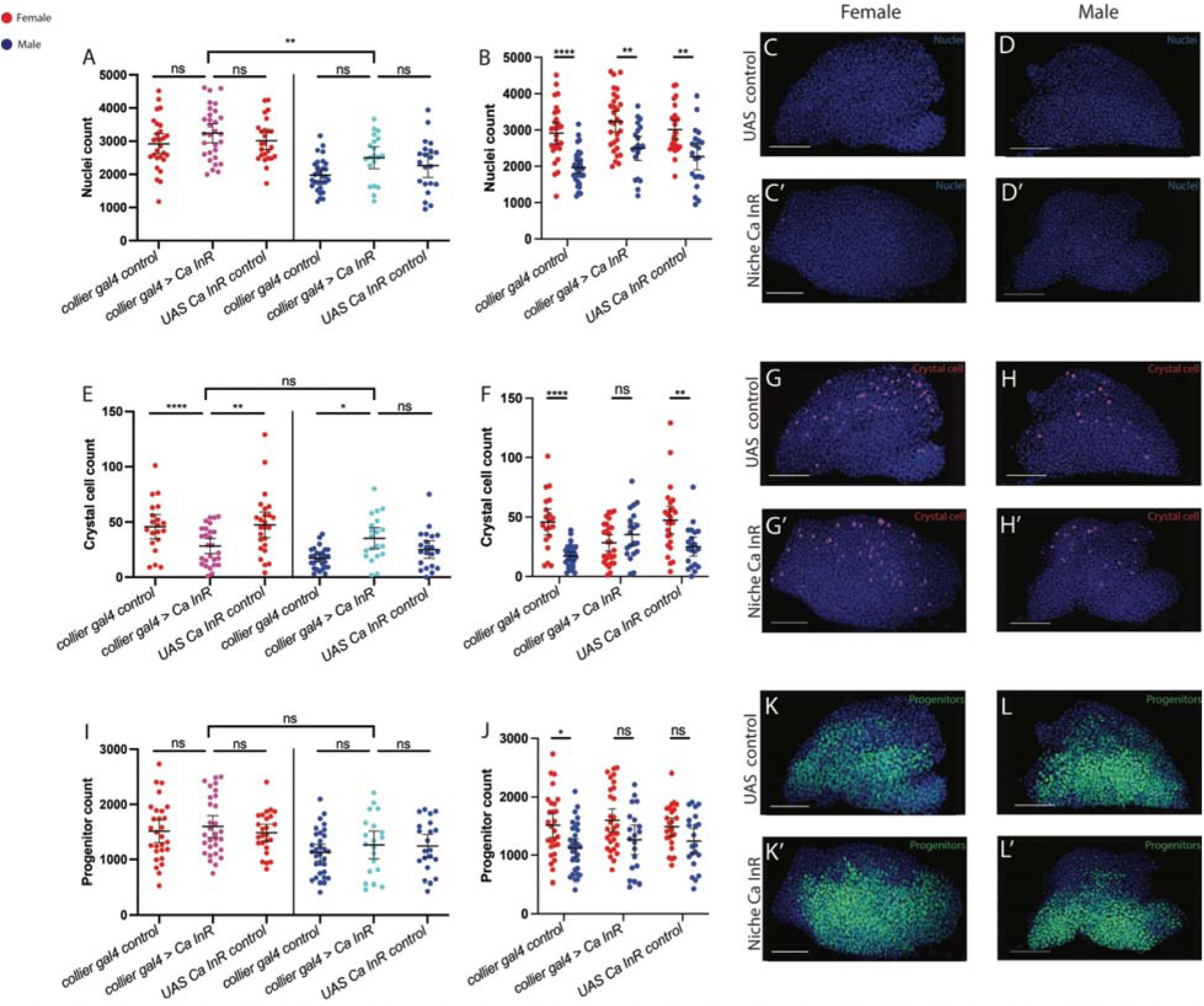
Constitutively activating insulin in the PSC. (A-B) Raw cell counts for total number of nuclei, stained with ToPro. Quantification using a two-way ANOVA, post-hoc Tukey’s test, for: *collier-Gal4* control females (n=28), *collier-Gal4>Ca InR* females (n=30), *UAS Ca InR* control females (n=25), *collier-Gal4* control males (n=31), *collier-Gal4>Ca InR* males (n=20), *UAS Ca InR* control males (n=21). The genotype:sex interaction constant was no significant (p=0.6852). (C) Representative image of *+> UAS Ca InR* control females stained with ToPro. (C’) Representative image of *collier-Gal4 > UAS Ca InR* female stained with ToPro. (D) Representative image of *+> UAS Ca InR* control male stained with ToPro, (D’) Representative image of *collier-Gal4 > UAS Ca InR* male stained with ToPro. (E-F) Raw cell counts for total number of crystal cells, stained with Hnt. Quantification using a two-way ANOVA, post-hoc Tukey’s test, for: *collier-Gal4* control females (n=20), *collier-Gal4>Ca InR* females (n=27), *UAS Ca InR* females (n=25), *collier-Gal4* control males (n=30), *collier-Gal4>Ca InR* males (n=20), *UAS Ca InR* control males (n= 21). The genotype:sex interaction constant was significant (p<0.0001). (G) Representative image of *+> UAS Ca InR* control females stained with Hnt. (G’) Representative image of *collier-Gal4 > UAS Ca InR* female stained with Hnt. (H) Representative image of *+> UAS Ca InR* control male stained with Hnt (H’) Representative image of *collier-Gal4 > UAS Ca InR* male stained with Hnt. (I-J) Raw cell counts for total number of progenitors, expressing GFP in domeMESO+ cells. Quantification using a two-way ANOVA, post-hoc Tukey’s test, for:*collier-Gal4* control females (n=28), *collier-Gal4>Ca InR* females (n=30), *UAS Ca InR* control females (n=25), *collier-Gal4* control males (n=31), *collier-Gal4>Ca InR* males (n=20), *UAS Ca InR* control males (n=21). The genotype:sex interaction constant was not significant (p=0.7455). (K) Representative image of *+> UAS Ca InR* control females expressing GFP in domeMESO+ cells. (K’) Representative image of *collier-Gal4 > UAS Ca InR* female expressing GFP in domeMESO+ cells. (L) Representative image of *+> UAS Ca InR* control male expressing GFP in domeMESO+ cells. (L’) Representative image of *collier-Gal4 > UAS Ca InR* male expressing GFP in domeMESO+ cells.**** indicates P<0.0001, *** indicates P<0.001, ** indicates P<0.01, * indicates P<0.05, ns (non-significant) indicates P>0.05. Error bar indicates 95% CI.

We found that PSC-specific reduction in insulin signaling by expression of InR-RNAi led to an increase in the number of cells in male lymph glands, but not female lymph glands (Figure 7A-D). However, the magnitude of the effect was not statistically significant between the sexes (sex:genotype p= 0.0931; two-way ANOVA; Table 2). In comparison, crystal cell numbers exhibited a more dramatic effect, as their number was greater in experimental group males in which InR was knocked down in the PSC than in females of comparable genotype (Figure 7E-H). Importantly, we observed a significant sex:genotype interaction and the magnitude of the effect was greater in males than in females, which eliminated the sex difference in this trait (sex:genotype p< 0.0001 Table 2). Analysis of progenitor numbers following InR knockdown in the PSC did not provide statistically significant support for changes in sex differences in this trait (Figure 7I-L) (sex:genotype p= 0.5789; Table 2).

**Table 2.** Interaction constants for insulin modulation. Sex:Genotype interaction constants derived from two-way ANOVA, post-hoc Tukey’s test performed on data from Figures 7-8(see methods). Significant interaction constant highlighted in yellow(p<0.05).

|  | Nuclei | Crystal cells | Progenitors |
| --- | --- | --- | --- |
| InR RNAi (Fig 7) | 0.0931 | <0.0001 | 0.5789 |
| Ca InR (Fig 8) | 0.6852 | <0.0001 | 0.7455 |

Analysis of PSC-specific insulin pathway activation via expression of Ca-InR revealed a similar trend. PSC-specific expression of Ca-InR did not greatly alter the number of cells in either male or female lymph glands and consequently there was no statistically significant impact on sex differences in organ size between females and males (Figure 8A-D) (sex:genotype p= 0.6852; Table 2). In contrast, crystal cell numbers were decreased by PSC specific activation of insulin signaling in females, which resulted in a statistically significant impact leading to an elimination of sex differences (Figure 8E-H) (sex:genotype p<0.0001; Table 2). Similar to overall lymph gland size, progenitor numbers were not impacted by PSC specific insulin pathway activation in either males or females and consequently there did not appear to be a significant reduction in sex differences (Figure 8I-L) (sex:genotype p=0.7455; Table 2).

Based on the two sets of experiments, utilizing downregulation or hyperactivation of the insulin receptor in the PSC we do not find conclusive support for a function of the insulin pathway in the PSC in mediating sex differences in overall lymph gland size or progenitor numbers. However, there was robust statistical support for the finding that the insulin pathway in the PSC mediates sex differences in crystal cell number.

### Sex-differences are observed in the cellular immune response following infection

Previous studies in adult *Drosophila* identified sex-specific differences in the immune response to infection that led to variation in survival rates. Intriguingly, these differences appeared to vary substantially based on the strain of bacteria used for infection [31]. We analysed the cellular immune response in males and females of wild-type (Canton-S) exposed to two different strains of bacteria that are known to activate the immune response in flies, *Escherichia coli (E. coli)* and *Pectobacterium carotovorum* (*P. carotovorum*; previously known as *Erwinia carotovora carotovora*, or *Ecc15)* [76,77](see methods). These strains were chosen because they have been shown to act in a sex-specific manner to control behaviour in adult flies [31,77,78,79]. Infection by feeding with *E.coli* showed a sex-specific increase in organ size and crystal cell count in females, but not in males (Figure 9A-C, D-F, Table 3). In comparison, neither sex showed an increase in progenitor count (Figure 9G-I, Table 3), while both sexes showed an increase in plasmatocyte count upon infection (Figure 9J-L, Table 3).

**Figure 9.**
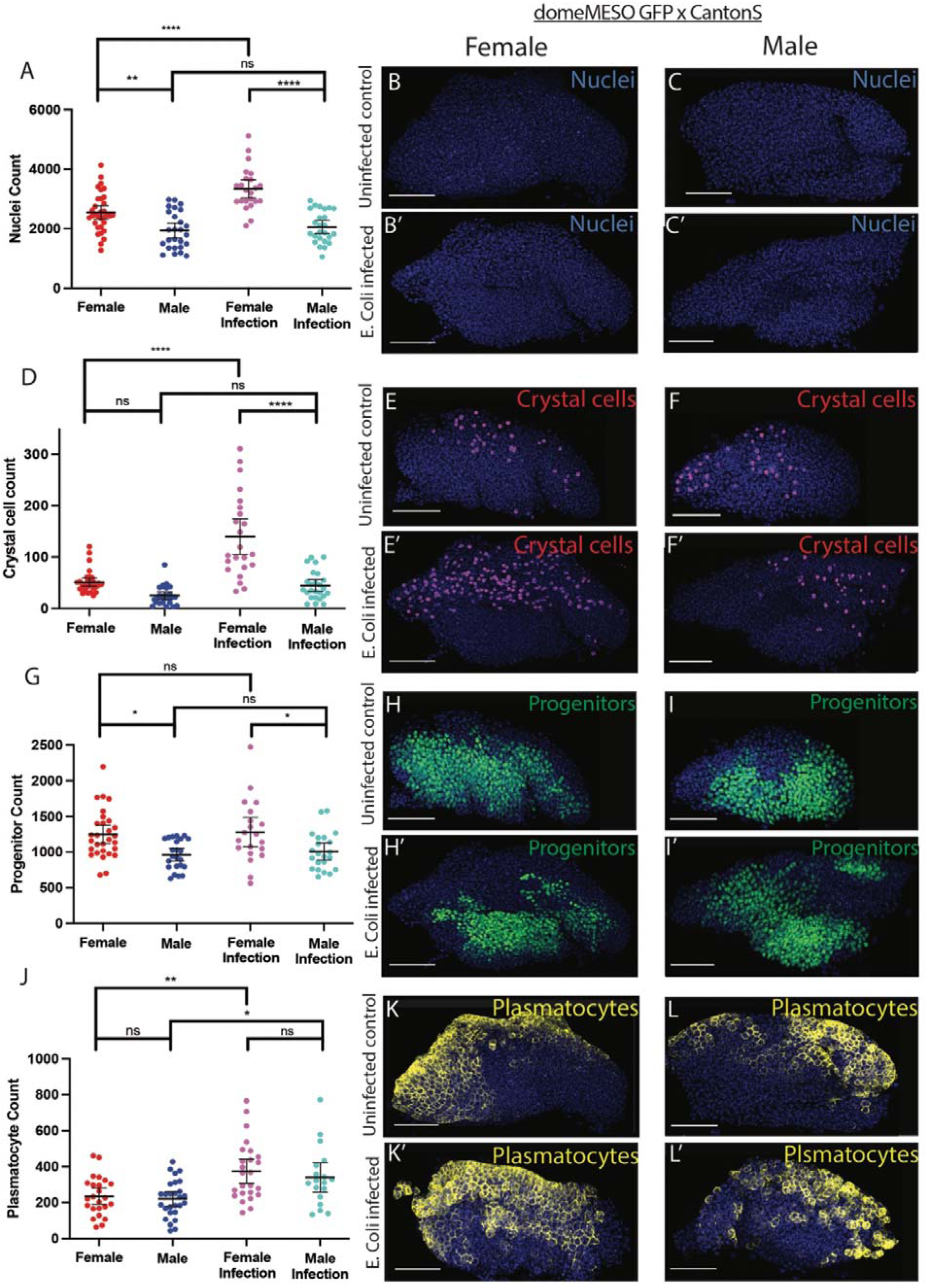
Infection with E. Coli. (A) Raw cell counts for total number of nuclei, stained with ToPro, in *domeMESO GFP x CantonS* flies both uninfected (female n=33, male n=26) and infected with *E.coli* bacteria (female n=23, male n=25). Sex:treatment interaction constant was significant (p=0.0076). (B, B’) Representative image of an uninfected (B) and infected (B’) female lymph gland stained with ToPro. (C, C’) Representative image of uninfected (C) and infected (C’) male lymph gland stained with ToPro. (D) Raw cell counts for total number of crystal cells in *domeMESO GFP x CantonS* flies both uninfected (female n=33, male n=26) and infected with *E.coli* bacteria (female n=23, male n=25). Sex:treatment interaction constant was significant (p<0.0001). (E, E’) Representative image of an uninfected (E) and infected (E’) female lymph gland stained with Hnt. (F, F’) Representative image of an uninfected (F) and infected (F’) male lymph gland stained with Hnt. (G) Raw cell counts for total number of progenitors, expressing GFP in domeMESO+ cells, in *domeMESO GFP x CantonS* flies both uninfected (female n=29, male n=23) and infected with *E.coli* bacteria (female n=20, male n=22). Sex:treatment interaction constant was not significant (p=0.9293). (H, H’) Representative image of an uninfected (H) and infected (H’) female lymph gland expressing GFP in domeMESO+ cells. (I, I’) Representative image of an uninfected male (I) and infected (I’) lymph gland expressing GFP in domeMESO+ cells. (J) Raw cell counts for total number of plasmatocytes, stained with P1, in *domeMESO GFP x CantonS* flies both uninfected (female n=24, male n=27) and infected with *E.coli* bacteria (female n=25, male n=18). Sex:treatment interaction constant was not significant (p=0.7160). (K, K’) Representative image of an uninfected (K) and infected (K’) female lymph gland stained with P1. (L, L’) Representative image of an uninfected (L) and infected (L’) male lymph gland stained with P1. All quantification done using a two-way ANOVA, post-hoc Tukey’s test. **** indicates P<0.0001, *** indicates P<0.001, ** indicates P<0.01, * indicates P<0.05, ns (non-significant) indicates P>0.05. Error bar indicates 95% CI.

**Table 3.** Interaction constants for infection. Sex:treatment interaction constants derived from two-way ANOVA, post-hoc Tukey’s test performed on data from Figures 9-10(see methods). Significant interaction constant highlighted in yellow(p<0.05).

|  | Nuclei | Crystal cells | Progenitors | Plasmatocytes |
| --- | --- | --- | --- | --- |
| *E. coli* | 0.0076 | <0.0001 | 0.9293 | 0.7160 |
| *P. carotovorum* | 0.3380 | 0.1627 | 0.0981 | 0.7884 |

Infection with *P. carotovorum* resulted in a significant increase in crystal cell numbers in females but not in males but the lack of a sex:treatment interaction meant that the observed effect of infection did not differ in magnitude between males and females (Figure 10D-F, table 3). In addition, overall organ size, progenitor numbers, and plasmatocyte numbers were not impacted by infection in either males or females (Figure 10A-C, G-I, J-L, Table 3). Taken together these data show that for *E. coli,* organ size and crystal cell production following infection was differentially regulated between male and female fly larvae. Moreover, analysis of infection with *P. carotovorum* showed that the magnitude or even the existence of sex-differences in the response to infection may vary depending on the bacterial strain used.

**Figure 10.**
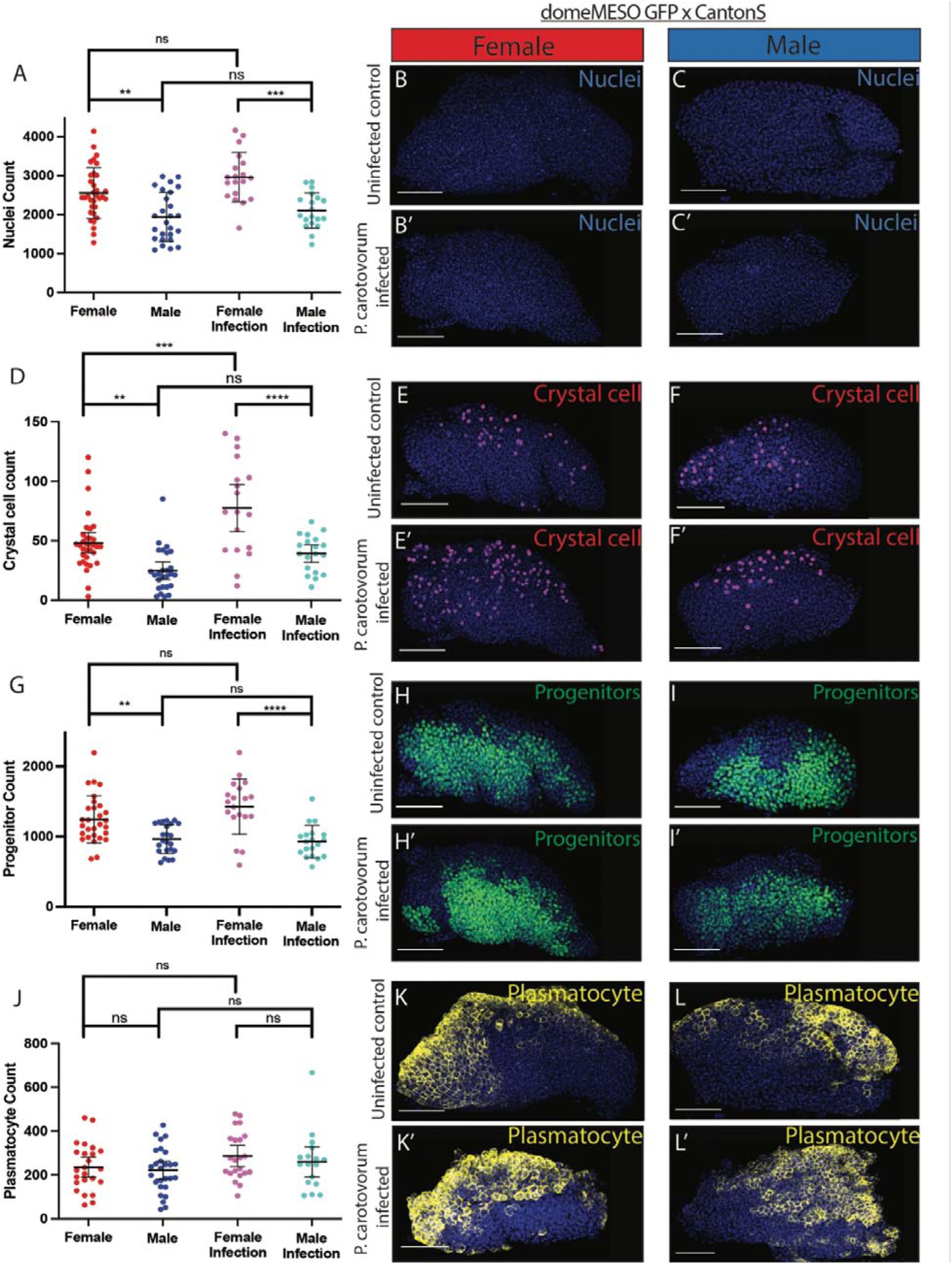
Infection with *P.carotovorum.* (A) Raw cell counts for total number of nuclei, stained with ToPro, in *domeMESO GFP x CantonS* flies both uninfected (female n=33, male n=26) and infected with *P. carotovorum* bacteria (female n=19, male n=19). Sex:treatment interaction constant was not significant (p=0.3380) (B, B’) Representative image of an uninfected (B) and infected (B’) female lymph gland stained with ToPro. (C) Representative image of uninfected (C) and infected (C’) male lymph gland stained with ToPro. (D) Raw cell counts for total number of crystal cells, stained with Hnt, in *domeMESO GFP x CantonS* flies both uninfected (female n=33, male n=26) and infected with *P.carotovorum* bacteria (female n=19, male n=19). Sex:treatment interaction constant was not significant (p=0.1627). (E, E’) Representative image of an uninfected (E) and infected (E’) female lymph gland stained with Hnt. (F, F’) Representative image of an uninfected (F) and infected (F’) male lymph gland stained with Hnt. (G)Raw cell counts for total number of progenitors, expressing GFP in domeMESO+ cells, in *domeMESO GFP x CantonS* flies both uninfected (female n=29, male n=23) and infected with *P. carotovorum* bacteria (female n=19, male n=19). Sex:treatment interaction constant was not significant (p=0.0981). (H, H’) Representative image of an uninfected (H) and infected (H’) female lymph gland expressing GFP in domeMESO+ cells. (I, I’) Representative image of an uninfected (I) and infected (I’) male lymph gland expressing GFP in domeMESO+ cells. (J) Raw cell counts for total number of plasmatocytes, stained with P1, in *domeMESO GFP x CantonS* flies both uninfected (female n=24, male n=27) and infected with *P. carotovorum* bacteria (female n=22, male n=17). Sex:treatment interaction constant was significant (p=0.7884). (K) Representative image of an uninfected female lymph gland stained with P1. (K’) Representative image of an infected female lymph gland stained with P1. (L) Representative image of an uninfected male lymph gland stained with P1. (L’) Representative image of an infected male lymph gland stained with P1. All quantification done using a one-way ANOVA, post-hoc Tukey’s test. **** indicates P<0.0001, *** indicates P<0.001, ** indicates P<0.01, * indicates P<0.05, ns (non-significant) indicates P>0.05. Error bar indicates 95% CI.

## Discussion

Our work establishes the *Drosophila* larva as a model for analysing sex-based differences in cellular immunity. In this regard we build upon an increasing body of evidence that multiple aspects of innate immunity in adult *Drosophila* exhibit striking sex-based variation [29–33,78–80]. We find that sex impacts the various cell types found in the lymph gland to different extents under both homeostatic or infection conditions, and that there are distinct mechanisms that control how sex differences are established in the various cell types. This supports a more nuanced and complex role for sex in regulating hematopoiesis beyond a simple generalised increased tissue size and cell number [76,81]. This idea is further supported by single-cell RNA-Seq that uncovered pervasive and wide ranging sex-differences at the level of gene expression across cell types in the lymph gland. Moreover, we found that the PSC, and more specifically insulin signaling from the PSC, mediated some sex differences but that there are other mechanisms in place since we were not able to mechanistically account for all the variance between males and females. Finally, in line with previous findings of sex-differences in the ability to survive infection [27,29–33] we find sex differences in the cellular immune response elicited by bacterial infection. Moreover, and consistent with previous findings [27,29–33], sex differences in the cellular immune response differed depending on the bacterial strain used for infection, indicating a complex relationship between sex, immunity, and infection. Taken together our work not only functions as an initial characterization of baseline sex-differences in fly hematopoiesis and in cellular immunity, but also identifies important areas for future exploration.

Our analysis of lymph gland single-cell RNA-Seq (scRNA-Seq) data shows significant differences in gene expression across multiple male and female cell types. These differences are not solely confined to genes involved in sex determination and X chromosome dosage compensation. We also observed sex-based expression differences in genes that have been implicated in hematopoiesis and blood cell function. Therefore, we hypothesize that some of the genes we found to be enriched in male or female cell types may be involved in generating the sex-specific hematopoietic phenotypes observed in lymph glands in this study. For example, we found that manipulating insulin signaling in the PSC in males, but not females, leads to changes in crystal cell differentiation. These results align well with the scRNA-Seq data, which shows that the insulin-like peptide dILP-6/Ilp6 is specifically enriched in male PSC cells and male crystal cells (Figure 2B). Future study is required to test if enrichment of dILP-6/Ilp6 expression in male PSC cells is an underlying mechanism contributing to this phenotype.

Although the existence of sex differences in innate immunity in *Drosophila* adults is well established [27,29,31], it is not yet standard practice to consider sex as a variable in studies that focus on the larval lymph gland. Based on our findings, we believe there is a substantial argument that in the future it should become a standard practice to consider sex as a key variable for all phenotypic characterization of the lymph gland.

This means that data should be reported separately for both sexes. Although some aspects of larval immunity, such as plasmatocyte numbers, may not vary greatly between males and females under normal conditions, differences might become apparent under specific experimental conditions. Our study did not aim to exhaustively characterize all sex-based differences in the lymph gland, which we suspect vary considerably based on the genetic background and experimental conditions. Rather our aim was to establish that sex differences are pervasive in the lymph gland and to provide some insight into the mechanisms that are responsible for these differences. We predict that as more researchers in the field separately report data for males and females for various lymph gland phenotypes, the number of documented sex differences will greatly expand.

Our analysis argues for an important role for the *Drosophila* lymph gland niche, the PSC, in at least a subset of sex differences. We did not find conclusive evidence for a role in sex differences for systemic signals that originate from the CNS and the fat body, or from local signals derived from the progenitors. These results suggest that juxtacrine signals or perhaps cell autonomous signals play an important mechanistic role in sex differences in hematopoiesis. However, this does not imply the absence of other known types of sex differences that have been demonstrated to exist in flies and that we did not assay for, such as differences in physiology or metabolism [81]. Instead our work assayed cell numbers, a parameter which is most relevant for immune function. As the repertoire of known sex differences in hematopoiesis grows, which we predict will be the case once separating data by sex becomes the norm in the field, it will be valuable to revisit the role of systemic signals as these are likely to play a part in at least some of these additional processes. Moreover, finding the mechanisms that mediate sex differences in progenitor numbers, which at present remain unknown, will be an important goal of future work.

In adult *Drosophila*, there are intriguing sex-specific behavioural responses to the presence of bacteria. For example, female flies are less likely to lay eggs in the presence of high levels of bacteria, seen as a protective mechanism designed to avoid egg laying in an environment inhospitable to them [77]. In the larval stage it is less clear what functions sex differences may have. It could be that their role is to set up future differences in the adult, and/or it could be that there is emphasis on preventing infection that could become chronic and harm egg production. The possibility of sex differences in larva impacting the adult immune response is supported by noting that a large portion of adult hemocytes derive from the larval lymph gland [82]. The potential importance of preventing infection in females during larval stages is highlighted by studies showing that infection in adult female flies impacts egg production and/or fecundity [83,84].

Taken as a whole, our work highlights the complex interactions between sex, genotype, and the environment in controlling *Drosophila* hematopoiesis and immunity. Our phenotypic analysis shows substantial variation across genotypes and environmental conditions in the proportion of different cell populations present in the lymph gland. In some experiments this did not allow us to obtain conclusive findings. However, this variability could represent important individual differences in life history, growth conditions, and environmental conditions between larvae. By accounting for sex as a factor that impacts lymph gland phenotype, it should be possible to reduce variation introduced by mixing data from males and females. This consideration will improve reproducibility and make future analysis more precise and meaningful.

## Materials and Methods

### Fly stocks and genetics

All *Drosophila* crosses were kept at 25 and sustained in vials containing a standard cornmeal fly food (recipe from Bloomington *Drosophila* Stock Center). All lymph gland progenitors were labelled using fluorescent markers *tep4-gal4; UAS-GFP or domeMESO GFP*. Control larvae in Figure 1 are categorized as *w^1118^*flies crossed to *tep4-gal4; UAS GFP*. Wildtype larvae in Figures 8-9 are categorized as CantonS flies crossed to *domeMESO GFP*.

Using the Gal4/UAS system [85], various tissues were feminized by overexpressing sex determination gene *transformer (*tra), a master regulator of female sexual identity and development [68–71,85]. The drivers used were *elav-gal4* (CNS) [86], *R4-gal4* (fat body) [87], *tep4-gal4* (lymph gland progenitors) [88], and *collier-gal4* (PSC cells) [89]. These drivers were crossed to *domeMESO GFP* for controls, and *UAS Tra^F^/CyoGFP; domeMESO GFP/TM6* for feminization experiments. A UAS control was also used by crossing *UAS Tra^F^/CyoGFP; domeMESO GFP/TM6* to *domeMESO GFP*.

In order to constitutively activate or knockdown insulin receptors in the lymph gland, *Ca-InR* (RRID: BDSC_8250) and *InR RNAi* (RRID: BDSC_51518) were crossed to *domeMESO GFP* as a control, and crossed to PSC cell driver *collier-gal4*.

### Larval lymph gland dissections

Wandering third instar larval lymph glands were dissected in ice-cold 1X Phosphate Buffer Saline (PBS), and then fixed in 4% paraformaldehyde (PFA) for 15 minutes. Samples were then washed in 0.1% PTX (1X PBS with 0.1% Triton X [Thermofisher Scientific, BP 151100]) twice for 5 minutes each, then blocked with 16% Neat Goat Serum (NGS) (ab7481, abcam) for 15 minutes, before being incubated in a primary antibody overnight at 4. The following day, samples were washed again twice in 0.1% PTX for 5 minutes each, blocked in 16% NGS for 15 minutes, before being incubated in a secondary antibody for two hours at room temperature. Samples were washed three times in 0.1% PTX for 5 minutes each, and mounted in VECTASHIELD (Vector Laboratories, H-1000, RRID: AB_2336789) with TOPRO3 iodide (ThermoFisher Scientific, T3605; 1:500) in glass bottom mounting dishes (MatTek Corporation, 35 mm, P35G-0-14-C).

### Immunohistochemistry

The following primary antibodies were used, and diluted in 0.1% PTX: mouse anti-hindsight (1:50, DSHB 1G9, RRID: AB_2617420), mouse anti-Antennapedia (1:25, DSHB 4C3, RRID: AB_528082). The following primary antibodies were used, and diluted in 0.1% Tween 20 (1X PBS with 0.1% Tween 20 [Fisher BioReagents, BP337-500]): mouse anti-P1 (1:100, anti-NimC1, a kind gift from Dr. Istvan Ando, Hungarian Academy of Sciences, Hungary).

The following secondary antibody was used, and diluted in 0.1% PTX (for Hnt and Antp staining) or 0.1% Tween 20 (for P1 staining): donkey anti-mouse Cy3 (1:400, Jackson Immunoresearch Laboratories, Code: 715-165-150, RRID: AB_2340813) and TOPRO3 iodide (1:800).

### Analysis of Single-cell RNA-Seq data

Partek Flow software was used for single-cell sequencing analysis. The previously published single-cell data set was used, which includes the same initial data processing parameters for read trimming, quality control, alignment of reads to the Drosophila melanogaster reference genome r6.22, normalization and low input filtration, and exclusion of ribosomal RNA genes [56].

Principal component (PC) analysis (PCA) was completed prior to graph-based clustering. Based on the Scree plot, the first 20 PCs were selected to use as the input for clustering and data visualization tasks. Graph-based clustering was first performed with 50 nearest neighbors (NNs) and resolution (res) of 0.25–0.75 giving 6–9 clusters, with 7 clusters (res=0.5-0.57) showing the most marked differences in gene expression between clusters. The data was visualized using UMAP with a local neighborhood size of 15, minimal distance of 0.1, Euclidean distance metric, and random generator seed of 0. We used established lymph gland marker genes to identify which cell type each of the graph-based clusters corresponds as done previously [53]. We determined sex by using the expression of two long non-coding RNAs (lncRNAs): lncRNA:roX1, lncRNA:roX2. Cells that are high for expression of both lncRNA:roX1 and lncRNA:roX2 were characterized as males, cells that were low for both were characterized as female, and cells that had high expression for only one of the lncRNAs were characterized as ambiguous and excluded from further analysis.

We used an ANOVA analysis to determine which genes were differentially expressed in males versus female cells. We set a threshold of 1.5-fold enrichment for genes to be considered differentially expressed and discarded genes which were enriched below that threshold. We also used a false-discovery rate (FDR) threshold of less than or equal to 0.0001, discarding any genes which were above that threshold. We took the remaining genes and thus had a list of differentially expressed genes (DEGs) for males and females (Supplementary file 1-Table 1). We then performed a similar analysis comparing male and female cells in each of the graph-based clusters to identify sex differences in specific lymph gland cell types (Supplementary file 1-Table 1).

We then compared the list of DEGs from each cluster to one another and to the DEGs for the lymph gland overall (Supplementary file 1-Table 4). We used the multiple list comparator from molbiotools to compare the lists and generate Venn diagrams to see which genes were uniquely over-expressed in each cluster. We then used g:Profiler’s g:GOSt tool [95] for gene-set enrichment analysis to find gene ontology terms, KEGG and WP pathways, or transcription factor binding sites associated with the DEGs in each cluster (Supplementary file 1-Table 2). We used these terms to highlight DEGs of particular interest due to their pattern of enrichment in specific male or female cell types, their involvement in blood cell development or disease, and their conservation across species (Supplementary file 1-Table 3).

### Infection

*Escherichia coli* (a kind gift from Dr. Bret Finlay, The University of British Columbia, Vancouver, Canada) and *Pectobactorium carotovorum* (a kind gift from Dr. Edan Foley, University of Alberta, Edmonton, Canada) were grown in LB media and incubated at 37 and 30 respectively overnight. Second instar larvae were picked out of food, washed in ddH2O and 70% ethanol, then starved on a polydimethylsiloxane pad for 2 hours before being placed into 4g of fly food with either 200 µl of PBS (control) or 200µL of *E.coli* or *P. carotovorum* resuspended in PBS. Vials were then placed back in 25 for 9 hours and dissected using the protocol described previously.

### Imaging and data analysis

All images were acquired on an Olympus FV1000 inverted confocal microscope, using a 40x lens. Image analysis was done using Olympus Fluoview (Ver.1.7c) and MATLAB scripts previously used in Khadiklar, 2017 [89]. Total number of nuclei, prohemocytes and plasmatocytes were determined using this Matlab script. Crystal cells and niche cells were counted manually using the cell counter plugin in ImageJ.

To correct for body size, average weight was taken for males and females of a genotype by collecting 10-12 third instar larvae, weighing them, and dividing total weight by number of larvae weighed. This weight, in milligrams, was then used to correct data points for body size (Figure 1F’-J’).

Statistics were performed using GraphPad Prism. P values were determined using a two-tailed unpaired t-test, a one way ANOVA with multiple comparisons (Tukey’s test), or a two-way ANOVA multiple comparison (Tukey’s Test). **** indicates P<0.0001, *** indicates P<0.001, ** indicates P<0.01, * indicates P<0.05, ns (non-significant) indicates P>0.05. The sample size and statistical method of each analysis were indicated in the figure legends.

A two-way ANOVA (Tukey’s test) was used to determine both p values in multiple comparisons, as well as an overall sex:genotype or sex:treatment interaction value. The multiple comparison significance are shown within the figure, with the exact p values and the interaction value shown in the figure legends.

## Supporting Information

**S1 File.** Table 1 Anova analysis was used to determine fold change in the number of reads for each gene, comparing female cells to male cells, as distinguished by expression of roX1 and rox2. We set a threshold of 1.5 fold change, and <=0.0001 for the FDR step-up, discarding any genes which did not meet both thresholds.

**S2 File.** Table 2 Using g:Profilers’s g:GOSt tool for gene-set enrichment analysis we analyzed lists of unique DEGs from Table 4 (see highlighted columns) to find gene ontology terms, pathways, and transcription factor targets that are enriched in each sex and cluster.

**S3 File.** Table 3 A selection of interesting sex-specific DEGs for further study. These genes were selected for being enriched in one cell type, being broadly conserved across species (including in humans in most cases), and having some connection to blood cell development or disease, or immune cell phenotypes. References are included for each gene detailing the known roles of these genes or their mammalian orthologs in blood cell development, disease, or phenotypes.

**S4 File.** Table 4 We used the multiple list comparator from molbiotools to compare DEG lists from Table 1 and determine which genes were uniquely overexpressed in each cluster. We ran this analysis both with and without the ‘overall’ category, which refers to the comparison of total male and female lymph gland cells.

