## Supplemental file 1 for "Sex differences in the regulation and function of cellular immunity in *Drosophila*"

| | A | B | C | D | E | F | G | H | I | J |
|---|---|---|---|---|---|---|---|---|---|---|
| 1 | **Female Genes overall** | | | | | **Male Genes Overalll** | | | | |
| 2 | **Gene** | **FDR** | **Fold change** | **LSMean** | | **Gene** | **FDR** | **Fold change** | **LSMean** | |
| 3 | lncRNA:CR40469 | 0.00E+00 | 25.66 | 381.49 | | lncRNA:roX1 | 0.00E+00 | ##### | 589.04 | |
| 4 | CG43133 | 0.00E+00 | 14.02 | 101.7 | | lncRNA:roX2 | 0.00E+00 | ##### | 311.00 | |
| 5 | CG32816 | 2.52E-181 | 4.43 | 10.245 | | msl-2 | 0.00E+00 | 3.90 | 16.18 | |
| 6 | CR43214 | 0.00E+00 | 3.92 | 16.865 | | CG32706 | ####### | 2.67 | 6.93 | |
| 7 | Cht2 | 3.16E-135 | 2.63 | 13.861 | | CG6999 | 1.88E-93 | 2.63 | 4.40 | |
| 8 | CG3038 | 1.13E-178 | 2.47 | 12.618 | | CG15739 | ####### | 2.56 | 9.43 | |
| 9 | CR43215 | 8.34E-77 | 2.45 | 2.8777 | | lncRNA:CR45979 | 8.87E-64 | 2.18 | 2.70 | |
| 10 | CR43211 | 0.00E+00 | 2.20 | 55.7 | | D2hgdh | ####### | 2.13 | 11.52 | |
| 11 | CG15784 | 1.59E-35 | 2.18 | 25.842 | | CG4586 | 1.44E-60 | 2.07 | 3.35 | |
| 12 | MCTS1 | 0.00E+00 | 2.16 | 145.67 | | comt | 2.25E-72 | 2.06 | 3.58 | |
| 13 | CG4301 | 1.41E-76 | 2.11 | 10.018 | | CG3568 | 1.96E-96 | 2.04 | 10.51 | |
| 14 | CG9733 | 6.28E-33 | 2.11 | 14.48 | | CG5273 | ####### | 2.03 | 8.93 | |
| 15 | Cpr5C | 2.87E-05 | 2.08 | 2.5977 | | CG10859 | 8.78E-47 | 2.03 | 3.06 | |
| 16 | Tsp3A | 1.77E-114 | 2.01 | 10.893 | | jv | 1.59E-25 | 1.98 | 6.33 | |
| 17 | NimC1 | 2.36E-29 | 2.01 | 16.193 | | CG16957 | 2.88E-34 | 1.84 | 2.90 | |
| 18 | CG14434 | 0.00E+00 | 1.93 | 81.315 | | lncRNA:CR32582 | ####### | 1.82 | 18.83 | |
| 19 | CG32641 | 5.87E-60 | 1.90 | 7.3755 | | fs(1)Yb | 7.70E-51 | 1.81 | 1.94 | |
| 20 | CecB | 7.02E-143 | 1.87 | 386.9 | | CG17919 | 2.24E-12 | 1.79 | 3.62 | |
| 21 | sano | 4.07E-17 | 1.85 | 6.8053 | | corolla | 5.62E-35 | 1.73 | 2.01 | |
| 22 | Sxl | 0.00E+00 | 1.79 | 67.863 | | CR18166 | 1.94E-43 | 1.72 | 3.26 | |
| 23 | CR43212 | 4.15E-85 | 1.79 | 14.163 | | CG7997 | ####### | 1.71 | 57.40 | |
| 24 | AnxB11 | 2.73E-279 | 1.78 | 91.913 | | CG11655 | 3.93E-65 | 1.71 | 7.39 | |
| 25 | Crg-1 | 1.98E-33 | 1.78 | 2.4806 | | lncRNA:CR45625 | 1.58E-49 | 1.68 | 5.81 | |
| 26 | dpr14 | 8.87E-30 | 1.75 | 3.7694 | | CG3176 | ####### | 1.68 | 20.13 | |
| 27 | CecA1 | 1.42E-115 | 1.73 | 199.71 | | CG15317 | 4.99E-64 | 1.68 | 10.23 | |
| 28 | CG34054 | 1.44E-41 | 1.72 | 22.984 | | AttA | 5.52E-08 | 1.66 | 7.24 | |
| 29 | Karl | 0.00E+00 | 1.66 | 453.78 | | CG42369 | 1.84E-56 | 1.66 | 47.76 | |
| 30 | wus | 4.81E-28 | 1.64 | 3.3782 | | CG33181 | 6.36E-25 | 1.64 | 4.20 | |

| | A | B | C | D | E | F | G | H | I | J |
|---|---|---|---|---|---|---|---|---|---|---|
| 31 | Arpc3B | 0.00E+00 | 1.64 | 300.2 | | sug | 4.54E-41 | 1.59 | 26.14 | |
| 32 | unc-13 | 1.83E-64 | 1.61 | 17.824 | | CG4239 | 1.04E-54 | 1.58 | 10.84 | |
| 33 | CG31821 | 2.05E-12 | 1.61 | 14.254 | | Su(Ste):CR42415 | 3.57E-21 | 1.58 | 1.58 | |
| 34 | mthl4 | 4.58E-24 | 1.60 | 6.6094 | | wde | 4.60E-91 | 1.56 | 35.11 | |
| 35 | DIP-alpha | 1.95E-31 | 1.59 | 6.8221 | | mof | 1.79E-38 | 1.56 | 6.63 | |
| 36 | CecC | 2.19E-98 | 1.59 | 1022.3 | | CG43867 | 1.00E-21 | 1.55 | 2.71 | |
| 37 | Tsp42Ed | 4.64E-06 | 1.58 | 4.105 | | CG5167 | 1.87E-87 | 1.53 | 28.91 | |
| 38 | prage | 2.10E-131 | 1.57 | 193.77 | | MtnA | 3.86E-26 | 1.53 | 390.56 | |
| 39 | CG33225 | 7.05E-25 | 1.57 | 27.969 | | aay | 2.45E-06 | 1.52 | 27.21 | |
| 40 | CG3587 | 1.38E-39 | 1.56 | 8.418 | | CkIIalpha-i3 | 1.54E-43 | 1.51 | 10.51 | |
| 41 | Rip11 | 4.06E-100 | 1.56 | 158.92 | | TM4SF | 5.63E-10 | 1.50 | 3.58 | |
| 42 | CG9981 | 5.42E-20 | 1.56 | 2.7021 | | | | | | |
| 43 | CG10804 | 9.43E-19 | 1.55 | 2.3367 | | | | | | |
| 44 | CecA2 | 4.25E-73 | 1.55 | 419.81 | | | | | | |
| 45 | CG4593 | 3.68E-177 | 1.55 | 71.598 | | | | | | |
| 46 | Sox15 | 1.10E-18 | 1.53 | 1.6546 | | | | | | |
| 47 | CG1840 | 8.05E-105 | 1.53 | 42.534 | | | | | | |
| 48 | lawc | 5.04E-120 | 1.52 | 59.021 | | | | | | |
| 49 | png | 1.68E-18 | 1.52 | 4.3765 | | | | | | |
| 50 | CG5004 | 1.12E-115 | 1.51 | 88.79 | | | | | | |
| 51 | CG3781 | 1.98E-109 | 1.51 | 36.792 | | | | | | |
| 52 | RhoGAP18B | 2.49E-182 | 1.51 | 143.34 | | | | | | |
| 53 | | | | | | | | | | |
| 54 | | | | | | | | | | |
| 55 | | | | | | | | | | |
| 56 | | | | | | | | | | |
| 57 | | | | | | | | | | |
| 58 | | | | | | | | | | |
| 59 | | | | | | | | | | |

| | K | L | M | N | O | P | Q | R | S | T |
|---|---|---|---|---|---|---|---|---|---|---|
| 1 | **Female Genes MZ** | | | | | **Male Genes MZ** | | | | |
| 2 | **Gene** | **FDR** | **Fold change** | **LSMean** | | **Gene** | **FDR** | **Fold change** | **LSMean** | |
| 3 | lncRNA:CR40469 | ####### | 36.88 | 340.59 | | lncRNA:roX1 | ####### | 596.90 | 596.90 | |
| 4 | CG43133 | ####### | 20.83 | 185.74 | | lncRNA:roX2 | ####### | 340.04 | 340.04 | |
| 5 | CR43214 | ####### | 3.95 | 17.40 | | msl-2 | ####### | 4.26 | 15.99 | |
| 6 | Cht2 | 8.06E-12 | 3.08 | 5.24 | | comt | 3.67E-28 | 2.73 | 3.66 | |
| 7 | CR43215 | 2.00E-26 | 2.65 | 3.08 | | CG15739 | 5.97E-22 | 2.41 | 6.07 | |
| 8 | CR43211 | ####### | 2.26 | 61.91 | | CG3176 | 2.89E-43 | 2.28 | 16.37 | |
| 9 | CG3038 | 3.95E-35 | 2.24 | 10.07 | | CG32706 | 1.03E-17 | 2.27 | 5.57 | |
| 10 | MCTS1 | ####### | 2.19 | 148.24 | | CG32625 | 2.68E-13 | 2.21 | 4.55 | |
| 11 | CG9733 | 2.75E-11 | 2.10 | 15.20 | | CG5273 | 2.93E-11 | 2.13 | 4.99 | |
| 12 | CG14434 | ####### | 2.05 | 99.87 | | CG11655 | 1.22E-16 | 2.13 | 5.60 | |
| 13 | CecB | 2.72E-46 | 1.97 | 419.45 | | CG6999 | 1.81E-10 | 2.08 | 3.00 | |
| 14 | CG15784 | 9.24E-08 | 1.96 | 11.55 | | lncRNA:CR32582 | 1.35E-35 | 2.06 | 16.03 | |
| 15 | CR43212 | 2.02E-39 | 1.94 | 16.33 | | lncRNA:CR4597 | 1.10E-08 | 1.87 | 2.33 | |
| 16 | Sxl | ####### | 1.94 | 61.22 | | CG43867 | 1.86E-08 | 1.86 | 2.64 | |
| 17 | CG31821 | 7.97E-13 | 1.91 | 14.98 | | CG16957 | 1.58E-08 | 1.82 | 3.07 | |
| 18 | CG32641 | 7.41E-17 | 1.88 | 6.57 | | CR18166 | 3.01E-12 | 1.81 | 3.10 | |
| 19 | lncRNA:CR43264 | 2.46E-06 | 1.87 | 5.79 | | CG4586 | 3.97E-07 | 1.79 | 2.34 | |
| 20 | wus | 1.27E-11 | 1.84 | 3.44 | | CG3568 | 9.47E-06 | 1.78 | 4.78 | |
| 21 | CecA1 | 8.94E-64 | 1.83 | 295.28 | | CG15317 | 1.10E-20 | 1.77 | 11.11 | |
| 22 | lncRNA:CR30009 | 4.44E-23 | 1.81 | 10.82 | | mof | 1.27E-15 | 1.77 | 7.01 | |
| 23 | AnxB11 | 1.39E-87 | 1.78 | 100.90 | | CkIIalpha-i3 | 2.86E-12 | 1.77 | 7.93 | |
| 24 | His2B:CG17949 | 2.93E-09 | 1.76 | 2.68 | | nkd | 6.66E-04 | 1.76 | 2.83 | |
| 25 | Tsp3A | 1.35E-19 | 1.74 | 8.76 | | CG1998 | 8.83E-06 | 1.76 | 3.08 | |
| 26 | CecC | 1.29E-45 | 1.72 | ##### | | CG5167 | 2.97E-19 | 1.72 | 21.20 | |
| 27 | CG3587 | 1.00E-15 | 1.71 | 8.08 | | D2hgdh | 4.97E-07 | 1.70 | 5.31 | |
| 28 | CG10804 | 2.33E-09 | 1.66 | 2.72 | | corolla | 2.37E-08 | 1.68 | 2.16 | |
| 29 | Fas3 | 2.10E-07 | 1.66 | 7.85 | | wde | 2.24E-41 | 1.67 | 44.62 | |
| 30 | Arpc3B | ####### | 1.65 | 240.51 | | CG7997 | 2.53E-36 | 1.65 | 38.94 | |

| | K | L | M | N | O | P | Q | R | S | T |
|---|---|---|---|---|---|---|---|---|---|---|
| 31 | Atf3 | 5.41E-64 | 1.64 | 58.45 | | GstT3 | 3.35E-19 | 1.65 | 14.81 | |
| 32 | Wsck | 2.93E-05 | 1.64 | 3.05 | | CG43759 | 6.20E-05 | 1.63 | 2.18 | |
| 33 | unc-13 | 2.35E-22 | 1.62 | 17.67 | | lncRNA:CR4562 | 1.61E-07 | 1.63 | 4.59 | |
| 34 | CG3781 | 1.23E-36 | 1.61 | 32.67 | | Hsp70Aa | 1.68E-09 | 1.59 | 42.11 | |
| 35 | CG12056 | 1.79E-11 | 1.61 | 9.63 | | CG6762 | 6.02E-11 | 1.59 | 10.61 | |
| 36 | Karl | ####### | 1.61 | 274.16 | | CG4239 | 3.13E-10 | 1.59 | 9.11 | |
| 37 | Chrac-16 | 3.12E-30 | 1.60 | 19.36 | | ewg | 5.52E-13 | 1.56 | 13.67 | |
| 38 | CG31689 | 1.09E-05 | 1.60 | 2.51 | | sdk | 2.74E-05 | 1.53 | 2.59 | |
| 39 | CecA2 | 3.74E-54 | 1.60 | 815.30 | | CG32817 | 2.60E-06 | 1.50 | 2.44 | |
| 40 | lncRNA:CR45517 | 1.81E-08 | 1.60 | 4.14 | | | | | | |
| 41 | DIP-alpha | 3.06E-11 | 1.60 | 6.84 | | | | | | |
| 42 | CG5004 | 1.72E-51 | 1.58 | 98.01 | | | | | | |
| 43 | CG33225 | 1.73E-26 | 1.58 | 73.29 | | | | | | |
| 44 | Crg-1 | 8.60E-06 | 1.57 | 2.03 | | | | | | |
| 45 | CG4593 | 1.96E-58 | 1.57 | 72.43 | | | | | | |
| 46 | CR43217 | 4.89E-07 | 1.56 | 2.44 | | | | | | |
| 47 | CG17982 | 2.82E-35 | 1.56 | 38.32 | | | | | | |
| 48 | CG11590 | 3.08E-56 | 1.55 | 60.98 | | | | | | |
| 49 | CG7322 | 9.06E-25 | 1.55 | 25.78 | | | | | | |
| 50 | CG34331 | 9.85E-54 | 1.55 | 63.85 | | | | | | |
| 51 | hpRNA:CR32207 | 1.99E-14 | 1.55 | 17.62 | | | | | | |
| 52 | lncRNA:CR42868 | 1.05E-14 | 1.53 | 9.64 | | | | | | |
| 53 | RNaseP:RNA | 1.70E-06 | 1.53 | 2.23 | | | | | | |
| 54 | Dab | 2.24E-06 | 1.53 | 4.46 | | | | | | |
| 55 | Gclc | 3.62E-34 | 1.52 | 72.53 | | | | | | |
| 56 | CG2444 | 6.36E-05 | 1.52 | 445.41 | | | | | | |
| 57 | rdgB | 1.22E-20 | 1.51 | 15.58 | | | | | | |
| 58 | CG17109 | 1.58E-04 | 1.51 | 7.16 | | | | | | |
| 59 | Aven | 4.82E-08 | 1.50 | 7.00 | | | | | | |

| | U | V | W | X | Y | Z | AA | AB | AC | AD |
|---|---|---|---|---|---|---|---|---|---|---|
| 1 | **Female Genes CCs** | | | | | **Male Genes CCs** | | | | |
| 2 | **Gene** | **FDR** | **Fold change** | **LSMean** | | **Gene** | **FDR** | **Fold change** | **LSMean** | |
| 3 | lncRNA:CF | ####### | 25.78 | 276.85 | | lncRNA:roX1 | ####### | 555.08 | 555.08 | |
| 4 | CG43133 | 1.00E+00 | 5.33 | 17.22 | | lncRNA:roX2 | ####### | 205.59 | 205.59 | |
| 5 | CG32816 | 1.00E+00 | 3.65 | 7.15 | | jv | 1.26E-05 | 7.45 | 18.19 | |
| 6 | CG17778 | 5.18E-01 | 2.97 | 5.49 | | msl-2 | 7.19E-49 | 6.18 | 30.48 | |
| 7 | Gasp | 2.23E-03 | 2.72 | 3.01 | | CG15739 | 5.54E-86 | 5.11 | 88.87 | |
| 8 | MCTS1 | 1.42E-12 | 2.22 | 136.44 | | lncRNA:CR459 | 1.62E-10 | 3.71 | 4.14 | |
| 9 | CG14434 | 4.08E-02 | 2.16 | 64.63 | | CG6999 | 9.51E-04 | 3.30 | 3.62 | |
| 10 | CG30148 | 5.83E-01 | 2.01 | 63.36 | | CG11318 | 7.55E-81 | 2.89 | 11.39 | |
| 11 | Karl | 5.55E-05 | 1.97 | 148.75 | | trol | 4.87E-08 | 2.74 | 20.88 | |
| 12 | CG10527 | 6.53E-01 | 1.86 | 45.36 | | lncRNA:CR325 | 5.66E-14 | 2.18 | 24.46 | |
| 13 | alpha-Est4 | 1.00E+00 | 1.82 | 3.81 | | Ilp6 | 8.55E-15 | 2.12 | 126.50 | |
| 14 | CG14629 | 7.31E-01 | 1.74 | 602.40 | | CG9119 | 6.40E-31 | 2.09 | 2819.06 | |
| 15 | CG4593 | 1.69E-01 | 1.72 | 73.33 | | asRNA:CR4499 | 2.58E-22 | 2.05 | 5.85 | |
| 16 | CG14270 | 9.01E-01 | 1.71 | 32.62 | | E23 | 2.50E-06 | 2.01 | 4.28 | |
| 17 | lncRNA:CF | 1.00E+00 | 1.68 | 4.96 | | snRNA:U6:96A | 7.97E-04 | 2.01 | 2.01 | |
| 18 | CrebB | 9.98E-01 | 1.66 | 38.88 | | D2hgdh | 5.65E-13 | 1.82 | 25.84 | |
| 19 | Sxl | 3.37E-02 | 1.65 | 82.29 | | CG34232 | 2.09E-06 | 1.82 | 25.58 | |
| 20 | CG11590 | 2.59E-01 | 1.58 | 51.98 | | Jheh3 | 1.17E-26 | 1.78 | 4.77 | |
| 21 | Arpc3B | 1.00E+00 | 1.57 | 212.06 | | mthl14 | 1.51E-09 | 1.76 | 2.23 | |
| 22 | Galphai | 1.00E+00 | 1.56 | 63.82 | | Hk | 7.70E-08 | 1.71 | 4.39 | |
| 23 | Tep1 | 1.00E+00 | 1.56 | 152.46 | | CG2017 | 2.34E-05 | 1.70 | 10.88 | |
| 24 | CG7453 | 8.17E-01 | 1.54 | 54.24 | | lncRNA:CR430 | 4.05E-04 | 1.70 | 17.93 | |
| 25 | CG3939 | 2.21E-11 | 1.51 | 72.77 | | Tsp42Er | 4.71E-46 | 1.67 | 15.87 | |
| 26 | | | | | | CG5418 | 8.46E-52 | 1.66 | 193.89 | |
| 27 | | | | | | lncRNA:CR448 | 3.95E-31 | 1.66 | 2.21 | |
| 28 | | | | | | Gp150 | 3.95E-04 | 1.65 | 10.71 | |
| 29 | | | | | | Vav | 7.42E-08 | 1.62 | 21.31 | |
| 30 | | | | | | asRNA:CR4612 | 1.99E-04 | 1.62 | 2.75 | |

| | U | V | W | X | Y | Z | AA | AB | AC | AD |
|---|---|---|---|---|---|---|---|---|---|---|
| 31 | | | | | | CG7997 | 1.16E-04 | 1.60 | 38.02 | |
| 32 | | | | | | AsnS | 5.03E-07 | 1.59 | 41.05 | |
| 33 | | | | | | CG34120 | 3.08E-12 | 1.59 | 2.34 | |
| 34 | | | | | | PCNA2 | 1.47E-05 | 1.57 | 2.71 | |
| 35 | | | | | | CG11576 | 7.67E-19 | 1.57 | 24.41 | |
| 36 | | | | | | lncRNA:CR443 | 8.54E-11 | 1.56 | 1.92 | |
| 37 | | | | | | CG42240 | 4.56E-06 | 1.55 | 7.76 | |
| 38 | | | | | | mwh | 1.79E-33 | 1.53 | 4.94 | |
| 39 | | | | | | lncRNA:CR447 | 1.44E-10 | 1.53 | 2.24 | |
| 40 | | | | | | CG6805 | 1.17E-07 | 1.52 | 66.02 | |
| 41 | | | | | | CG9008 | 7.10E-04 | 1.51 | 4.50 | |
| 42 | | | | | | | | | | |
| 43 | | | | | | | | | | |
| 44 | | | | | | | | | | |
| 45 | | | | | | | | | | |
| 46 | | | | | | | | | | |
| 47 | | | | | | | | | | |
| 48 | | | | | | | | | | |
| 49 | | | | | | | | | | |
| 50 | | | | | | | | | | |
| 51 | | | | | | | | | | |
| 52 | | | | | | | | | | |
| 53 | | | | | | | | | | |
| 54 | | | | | | | | | | |
| 55 | | | | | | | | | | |
| 56 | | | | | | | | | | |
| 57 | | | | | | | | | | |
| 58 | | | | | | | | | | |
| 59 | | | | | | | | | | |

| | AE | AF | AG | AH | AI | AJ | AK | AL |
|---|---|---|---|---|---|---|---|---|
| 1 | **Female Genes PSC** | | | | | **Male Genes PSC** | | |
| 2 | **Gene** | **FDR** | **Fold change** | **LSMean** | | **Gene** | **FDR** | **Fold change** |
| 3 | lncRNA:CR40469 | ######## | 160.00 | 1036.74 | | lncRNA:roX1 | 1.84E-295 | 757.50 |
| 4 | CG6287 | 9.72E-05 | 1.78 | 797.79 | | lncRNA:roX2 | 1.38E-303 | 729.46 |
| 5 | mthl7 | 1.93E-09 | 1.55 | 259.86 | | Lcp65Ab1 | 3.42E-07 | 328.85 |
| 6 | CG15784 | 6.51E-12 | 2.22 | 234.56 | | Cpr47Ec | 2.58E-06 | 227.69 |
| 7 | Karl | 5.55E-05 | 1.52 | 212.68 | | CG34227 | 8.84E-06 | 166.00 |
| 8 | NimB5 | 5.21E-08 | 1.54 | 183.97 | | Lcp65Ab2 | 1.26E-06 | 153.09 |
| 9 | CG3939 | 2.21E-11 | 2.04 | 177.26 | | CG13403 | 1.22E-09 | 114.35 |
| 10 | Cpr5C | 0.00E+00 | 6.24 | 150.59 | | dpy | 3.89E-11 | 67.00 |
| 11 | ImpL2 | 6.87E-07 | 1.82 | 141.46 | | Ets98B | 1.20E-14 | 45.48 |
| 12 | CG10508 | 3.26E-09 | 1.53 | 130.26 | | fj | 1.18E-16 | 41.17 |
| 13 | Vps2 | 1.06E-04 | 1.75 | 126.61 | | SKIP | 4.65E-17 | 29.50 |
| 14 | MCTS1 | 1.42E-12 | 2.71 | 119.63 | | slf | 2.93E-07 | 26.83 |
| 15 | CR43211 | 4.07E-04 | 2.20 | 91.09 | | CG9449 | 2.72E-05 | 23.62 |
| 16 | Hers | 6.95E-04 | 1.83 | 90.96 | | CG34398 | 8.91E-05 | 15.35 |
| 17 | CG13369 | 2.14E-05 | 2.02 | 90.33 | | pk | 4.59E-13 | 10.98 |
| 18 | CG12112 | 2.10E-04 | 1.77 | 89.77 | | ovo | 2.14E-05 | 7.61 |
| 19 | TkR99D | ######## | 2.32 | 83.48 | | lncRNA:CR43432 | 4.87E-05 | 5.42 |
| 20 | CG43759 | 2.23E-15 | 1.67 | 66.29 | | mirr | 4.62E-06 | 4.48 |
| 21 | CG9220 | 2.19E-24 | 1.66 | 39.03 | | CG8526 | 2.31E-05 | 3.87 |
| 22 | CG14855 | 1.78E-24 | 1.90 | 38.67 | | CG46339 | 1.48E-11 | 3.33 |
| 23 | hlk | 9.73E-09 | 1.58 | 37.03 | | CG34370 | 3.05E-04 | 3.26 |
| 24 | snRNA:U2:14B | 7.98E-06 | 2.33 | 34.09 | | NetA | 1.42E-43 | 3.05 |
| 25 | CG15365 | 3.08E-16 | 1.56 | 29.77 | | Picot | 1.90E-04 | 3.00 |
| 26 | comm | 1.50E-06 | 1.64 | 20.75 | | axed | 9.35E-07 | 2.95 |
| 27 | lncRNA:CR44933 | 2.72E-22 | 1.53 | 20.26 | | NetB | 1.23E-09 | 2.90 |
| 28 | CG8369 | 7.17E-09 | 1.72 | 15.85 | | Ilp6 | 6.10E-07 | 2.77 |
| 29 | Ace | 1.53E-10 | 1.75 | 14.94 | | CG8547 | 5.65E-19 | 2.69 |
| 30 | CCKLR-17D3 | 2.21E-11 | 1.61 | 14.16 | | Hsp70Bb | 6.77E-05 | 2.67 |

| | AE | AF | AG | AH | AI | AJ | AK | AL |
|---|---|---|---|---|---|---|---|---|
| 31 | Idgf4 | 1.01E-23 | 3.51 | 13.60 | | Ca-alpha1D | 4.88E-07 | 2.67 |
| 32 | Dl | 8.10E-08 | 4.20 | 12.82 | | Npc2a | 5.96E-09 | 2.60 |
| 33 | asRNA:CR45912 | 2.40E-45 | 4.58 | 11.42 | | CG1113 | 2.30E-04 | 2.58 |
| 34 | comm2 | 1.61E-06 | 1.77 | 8.73 | | rdgA | 9.57E-35 | 2.48 |
| 35 | CG31183 | 2.14E-05 | 1.92 | 6.68 | | CG32280 | 3.15E-05 | 2.40 |
| 36 | lncRNA:CR33218 | 2.62E-04 | 5.03 | 5.03 | | lncRNA:CR45959 | 5.38E-06 | 2.39 |
| 37 | CG34002 | 9.26E-07 | 2.42 | 4.97 | | bark | 2.03E-09 | 2.30 |
| 38 | CG14516 | 1.96E-12 | 3.00 | 4.94 | | CG13284 | 8.48E-64 | 2.17 |
| 39 | form3 | 5.28E-04 | 2.55 | 4.69 | | CG9095 | 2.83E-07 | 2.16 |
| 40 | Septin4 | 9.72E-05 | 4.01 | 4.01 | | CG2781 | 2.99E-33 | 2.08 |
| 41 | CG17321 | 7.43E-07 | 3.27 | 3.27 | | ko | 1.04E-23 | 1.96 |
| 42 | CG17121 | 5.33E-07 | 3.12 | 3.12 | | GILT2 | 1.62E-31 | 1.96 |
| 43 | Dh31-R | 3.41E-04 | 3.05 | 3.05 | | CG11134 | 9.59E-06 | 1.66 |
| 44 | | | | | | Ac13E | 3.00E-04 | 1.66 |
| 45 | | | | | | Pgant2 | 2.11E-26 | 1.54 |
| 46 | | | | | | | | |
| 47 | | | | | | | | |
| 48 | | | | | | | | |
| 49 | | | | | | | | |
| 50 | | | | | | | | |
| 51 | | | | | | | | |
| 52 | | | | | | | | |
| 53 | | | | | | | | |
| 54 | | | | | | | | |
| 55 | | | | | | | | |
| 56 | | | | | | | | |
| 57 | | | | | | | | |
| 58 | | | | | | | | |
| 59 | | | | | | | | |

| | AM | AN | AO | AP | AQ | AR | AS | AT | AU |
|---|---|---|---|---|---|---|---|---|---|
| 1 | | | | | | | | | |
| 2 | **LSMean** | | | | | | | | |
| 3 | 757.50 | | | | | | | | |
| 4 | 729.46 | | | | | | | | |
| 5 | 328.85 | | | | | | | | |
| 6 | 227.69 | | | | | | | | |
| 7 | 166.00 | | | | | | | | |
| 8 | 153.09 | | | | | | | | |
| 9 | 114.35 | | | | | | | | |
| 10 | 67.00 | | | | | | | | |
| 11 | 45.48 | | | | | | | | |
| 12 | 41.17 | | | | | | | | |
| 13 | 29.50 | | | | | | | | |
| 14 | 26.83 | | | | | | | | |
| 15 | 23.62 | | | | | | | | |
| 16 | 15.35 | | | | | | | | |
| 17 | 10.98 | | | | | | | | |
| 18 | 7.61 | | | | | | | | |
| 19 | 5.42 | | | | | | | | |
| 20 | 4.48 | | | | | | | | |
| 21 | 5.58 | | | | | | | | |
| 22 | 7.82 | | | | | | | | |
| 23 | 5.23 | | | | | | | | |
| 24 | 7.14 | | | | | | | | |
| 25 | 7.67 | | | | | | | | |
| 26 | 10.87 | | | | | | | | |
| 27 | 8.78 | | | | | | | | |
| 28 | 144.20 | | | | | | | | |
| 29 | 13.68 | | | | | | | | |
| 30 | 10.15 | | | | | | | | |

| | AM | AN | AO | AP | AQ | AR | AS | AT | AU |
|---|---|---|---|---|---|---|---|---|---|
| 31 | 2.67 | | | | | | | | |
| 32 | 26.33 | | | | | | | | |
| 33 | 2.58 | | | | | | | | |
| 34 | 120.55 | | | | | | | | |
| 35 | 18.76 | | | | | | | | |
| 36 | 2.39 | | | | | | | | |
| 37 | 4.21 | | | | | | | | |
| 38 | 12.60 | | | | | | | | |
| 39 | 3.74 | | | | | | | | |
| 40 | 71.33 | | | | | | | | |
| 41 | 28.85 | | | | | | | | |
| 42 | 158.19 | | | | | | | | |
| 43 | 77.29 | | | | | | | | |
| 44 | 15.55 | | | | | | | | |
| 45 | 22.90 | | | | | | | | |
| 46 | | | | | | | | | |
| 47 | | | | | | | | | |
| 48 | | | | | | | | | |
| 49 | | | | | | | | | |
| 50 | | | | | | | | | |
| 51 | | | | | | | | | |
| 52 | | | | | | | | | |
| 53 | | | | | | | | | |
| 54 | | | | | | | | | |
| 55 | | | | | | | | | |
| 56 | | | | | | | | | |
| 57 | | | | | | | | | |
| 58 | | | | | | | | | |
| 59 | | | | | | | | | |

| | AV | AW | AX | AY | AZ | BA | BB | BC |
|---|---|---|---|---|---|---|---|---|
| 1 | | | | | | | | |
| 2 | | | | | | | | |
| 3 | | | | | | | | |
| 4 | | | | | | | | |
| 5 | | | | | | | | |
| 6 | | | | | | | | |
| 7 | | | | | | | | |
| 8 | | | | | | | | |
| 9 | | | | | | | | |
| 10 | | | | | | | | |
| 11 | | | | | | | | |
| 12 | | | | | | | | |
| 13 | | | | | | | | |
| 14 | | | | | | | | |
| 15 | | | | | | | | |
| 16 | | | | | | | | |
| 17 | | | | | | | | |
| 18 | | | | | | | | |
| 19 | | | | | | | | |
| 20 | | | | | | | | |
| 21 | | | | | | | | |
| 22 | | | | | | | | |
| 23 | | | | | | | | |
| 24 | | | | | | | | |
| 25 | | | | | | | | |
| 26 | | | | | | | | |
| 27 | | | | | | | | |
| 28 | | | | | | | | |
| 29 | | | | | | | | |
| 30 | | | | | | | | |
