## Supplemental file 2 for "Sex differences in the regulation and function of cellular immunity in *Drosophila*"

| Female Genes overall | | | | | Male Genes Overalll | | | |
|---|---|---|---|---|---|---|---|---|
| GO or pathway | ID | Adjusted p-value | Genes | | GO or pathway | ID | Adjusted p-value | Genes |
| flippase activity | GO:0140327 | 0.003349 | CG4301,CG9981 | | sex chromosome | GO:0000803 | 0.01370 | MSL-2,MOF |
| ATPase-coupled intramembrane lipid transporter | GO:0140326 | 0.049811 | CG4301,CG9981 | | X chromosome | GO:0000805 | 0.01370 | MSL-2,MOF |
| defense response to insect | GO:0002213 | 0.001438 | CECB,CECA1,CECA2 | | X chromosome located dosage compensation complex, transcription activating | GO:0016456 | 0.01370 | MSL-2,MOF |
| defense response to Gram-positive | GO:0050830 | 0.028312 | CECB,CECA1,CECC,CECA2 | | dosage compensation complex | GO:0046536 | 0.01370 | MSL-2,MOF |
| phospholipid-translocating ATPase complex | GO:1990531 | 0.005406 | CG4301,CG9981 | | MSL complex | GO:0072487 | 0.02550 | MSL-2,MOF |
| Toll and Imd signaling pathway | KEGG:04624 | 0.001696 | CECB,CECA1,CECC,CECA2 | | | | | |
| Intrahepatic cholestasis with episodic jaundice | HP:0006575 | 0.008293 | CG4301,CG9981 | | | | | |
| Dysphoria | HP:0033838 | 0.008293 | CG4301,CG9981 | | | | | |
| Abnormal liver function tests during pregnancy | HP:0200148 | 0.008293 | CG4301,CG9981 | | | | | |
| Increased serum bile acid concentration during pregnancy | HP:0200150 | 0.008293 | CG4301,CG9981 | | | | | |
| Asterixis | HP:0012164 | 0.016572 | CG4301,CG9981 | | | | | |

| | | | |
|---|---|---|---|
| Pruritus on foot | HP:0030900 | 0.016572 | CG4301,CG9981 |
| Abnormal speech discrimination | HP:0001963 | 0.027598 | CG4301,CG9981 |

| Female Genes MZ | | | | | Male Genes MZ | | | |
|---|---|---|---|---|---|---|---|---|
| GO or pathway | ID | Adjusted p-value | Genes | | GO or pathway | ID | Adjusted p-value | Genes |
| defense response to insect | GO:0002213 | 0.00099 | CECA2,CECB,CECA1 | | sex chromosome | GO:0000803 | 0.0137043 | MSL-2,MOF |
| defense response to Gram-positive bacterium | GO:0050830 | 0.01820 | CECC,CECA2,CECB,CECA1 | | X chromosome | GO:0000805 | 0.0137043 | MSL-2,MOF |
| antibacterial humoral response | GO:0019731 | 0.04017 | CECC,CECA2,CECB,CECA1 | | X chromosome located dosage compensation complex, transcription activating | GO:0016456 | 0.0137043 | MSL-2,MOF |
| Toll and Imd signaling pathway | KEGG:04624 | 0.00256 | CECC,CECA2,CECB,CECA1 | | dosage compensation complex | GO:0046536 | 0.0137043 | MSL-2,MOF |
| | | | | | MSL complex | GO:0072487 | 0.0255029 | MSL-2,MOF |

| Female Genes CCs | | | | | Male Genes CCs | | | |
|---|---|---|---|---|---|---|---|---|
| GO or pathway | ID | Adjusted p-value | Genes | | GO or pathway | ID | Adjusted p-value | Genes |
| No GO or pathway enrichment | | | | | Insect hormone biosynthesis | KEGG:00981 | 0.03545 | CG15739,JHEH3 |
| | | | | | Weak voice | HP:0001621 | 0.04914 | TROL,CG11576,CG6805 |

| | Female Genes PSC | | | | | Male Genes PSC | | | |
|---|---|---|---|---|---|---|---|---|---|
| | GO or pathway | ID | Adjusted p-value | Genes | | GO or pathway | ID | Adjusted p-value | Genes |
| | peptide receptor activity | GO:0001653 | 0.00249 | CCKLR-17D3,CG31183,TKR99D,DH31- | | regulation of axon guidance | GO:1902667 | 0.0235 | PK,NETA,NETB |
| | neuropeptide receptor activity | GO:0008188 | 0.03917 | CCKLR-17D3,TKR99D,DH31-R | | cell periphery | GO:0071944 | 0.0101 | LCP65AB1,CPR47EC,LCP65AB2 |
| | G protein-coupled receptor activity | GO:0004930 | 0.04506 | CCKLR-17D3,TKR99D,DH31-R,MTHL7 | | extracellular matrix | GO:0031012 | 0.0156 | LCP65AB1,CPR47EC,LCP65AB2,DPY,NETA,NETB |
| | Neuroactive ligand-receptor interaction | KEGG:04080 | 0.02683 | CCKLR-17D3,TKR99D,DH31-R | | external encapsulating structure | GO:0030312 | 0.0295 | LCP65AB1,CPR47EC,LCP65AB2 |
| | | | | | | extracellular region | GO:0005576 | 0.0343 | LCP65AB1,CG34227,LCP65AB2 [illegible] |
| | | | | | | Longevity regulating pathway - multiple species | KEGG:04213 | 0.0087 | ILP6,HSP70BB,AC13E |
