## Supplemental file 3 for "Sex differences in the regulation and function of cellular immunity in *Drosophila*"

| Gene | Enriched in | Summary | Reference |
|---|---|---|---|
| MCTS1 | female overall | Malignant T cell amplified sequence 1 is a translation reinitiation and ribosome recycling factor; mammalian ortholog is associated with Lymphoma and X-linked immunodeficiency | Proniak et al 2018; Bohlen et al 2023 |
| Ilp6 | male multiple zones (PSC and CC) | Insulin-like peptide involved in crystal cell development in response to CO2/O2 gaseous imblance; mammalian ortholog involved in thrombopoiesis | Cho et al. 2018; Chen et al. 2018 |
| TkR99D | female PSC | Tachykinin-like peptide receptor, a class of GPCRs that respond to neuropeptides; mammalian orthologs involved in hematopoiesis | Liu et al. 2007 |
| rdgA | male PSC | Diacylglycerol kinase involved in phospholipase C signaling; loss of the mammalian ortholog DGKZ results hyperactive T cell response | Zhong et al. 2002 |
| CecB | female MZ | Anti-microbial peptide involved in the immune response | Carboni et al. 2022 |
| CG5167 | male MZ | putative ortholog SCCPDH (saccharopine dehydrogenase) involved in glycolipid biosynthesis; expressed on alveolar macrophages | Patel et al. 2017 |
| Tep1 | female CC | Complement-like protein involved in the innate immune response; human ortholog CD109 is seen on activated platelets, T cells, and a subset of HSPCs | Dostalova et al. 2017; Batal et al. 2025 |
| CG15739 | male CC | phosphatase involved in vitamin B metabolism and juvenile hormone synthesis; human ortholog PDXP is expressed in RBCs | Gohla 2019 |
