## Supplemental file 4 for "Sex differences in the regulation and function of cellular immunity in *Drosophila*"

### Female DEGs by Zone

| PSC | CC | MZ | PSC and MZ | PSC and CCs | CC and MZ | PSC and CC and MZ |
|---|---|---|---|---|---|---|
| CG12112 | CG32816 | CecC | CG15784 | CG3939 | Arpc3B | Karl |
| Idgf4 | CG17778 | CecA2 | CR43211 | | CG11590 | MCTS1 |
| CG15365 | Gasp | CG2444 | | | CG14434 | lncRNA:CR40469 |
| CG14855 | CG30148 | CecB | | | CG43133 | |
| CG13369 | CG10527 | CecA1 | | | CG4593 | |
| form3 | alpha-Est4 | CG5004 | | | Sxl | |
| CCKLR-17D3 | CG14629 | AnxB11 | | | | |
| snRNA:U2:14B | CG14270 | Gclc | | | | |
| CG9220 | lncRNA:CR45530 | CG33225 | | | | |
| Hers | CrebB | CG34331 | | | | |
| CG31183 | Galphai | Atf3 | | | | |
| septin 4 | Tep1 | CG17982 | | | | |
| CG17321 | CG7453 | CG3781 | | | | |
| Dl | | CG7322 | | | | |
| Vps2 | | Chrac-16 | | | | |
| comm2 | | hpRNA:CR32207 | | | | |
| NimB5 | | unc-13 | | | | |
| lncRNA:CR33218 | | rdgB | | | | |
| CG10508 | | CR43212 | | | | |
| TkR99D | | CG31821 | | | | |
| CG14516 | | CG9733 | | | | |
| CG8369 | | lncRNA:CR42868 | | | | |
| lncRNA:CR44933 | | lncRNA:CR30009 | | | | |
| comm | | CG12056 | | | | |
| l(2)01289 | | Tsp3A | | | | |
| asRNA:CR45912 | | CG17109 | | | | |
| Ace | | CG3587 | | | | |
| Dh31-R | | Fas3 | | | | |
| CG17121 | | Aven | | | | |
| Cpr5C | | CG3038 | | | | |
| mthl7 | | CR43214 | | | | |
| CG34002 | | DIP-alpha | | | | |
| CG6287 | | CG32641 | | | | |
| ImpL2 | | lncRNA:CR43264 | | | | |
| CG43759 | | Dab | | | | |
| | | lncRNA:CR45517 | | | | |
| | | wus | | | | |
| | | Wsck | | | | |
| | | Cht2 | | | | |
| | | CG10804 | | | | |
| | | CG31689 | | | | |
| | | CR43217 | | | | |
| | | His2B:CG17949 | | | | |
| | | RNaseP:RNA | | | | |
| | | Crg-1 | | | | |
| | | CR43215 | | | | |

### Male DEGs by Zone

| PSC | CC | MZ | PSC and MZ | CC and MZ | PSC and CC | PSC and CC and MZ |
|---|---|---|---|---|---|---|
| Lcp65Ab1 | jv | comt | | CG15739 | Ilp6 | lncRNA:roX1 |
| Cpr47Ec | CG11318 | CG3176 | | CG6999 | | lncRNA:roX2 |
| CG34227 | trol | CG32706 | | CG7997 | | |
| Lcp65Ab2 | CG9119 | CG32625 | | D2hgdh | | |
| CG13403 | asRNA:CR44992 | CG5273 | | lncRNA:CR32582 | | |
| dpy | E23 | CG11655 | | lncRNA:CR45979 | | |
| Ets98B | snRNA:U6:96Ac | CG43867 | | msl-2 | | |
| fj | CG34232 | CG16957 | | | | |
| SKIP | Jheh3 | CR18166 | | | | |
| slf | mthl14 | CG4586 | | | | |
| CG9449 | Hk | CG3568 | | | | |
| CG34398 | CG2017 | CG15317 | | | | |
| pk | lncRNA:CR43051 | mof | | | | |
| ovo | Tsp42Er | CkIIalpha-i3 | | | | |
| lncRNA:CR43432 | CG5418 | nkd | | | | |
| mirr | lncRNA:CR44895 | CG1998 | | | | |
| CG8526 | Gp150 | CG5167 | | | | |
| CG46339 | Vav | corolla | | | | |
| CG34370 | asRNA:CR46128 | wde | | | | |
| NetA | AsnS | GstT3 | | | | |
| Picot | CG34120 | CG43759 | | | | |
| axed | PCNA2 | CG43051 | | | | |
| NetB | CG11576 | Hsp70Aa | | | | |
| CG8547 | lncRNA:CR44380 | CG6762 | | | | |
| Hsp70Bb | CG42240 | CG4239 | | | | |
| Ca-alpha1D | mwh | ewg | | | | |
| Npc2a | lncRNA:CR44757 | sdk | | | | |
| CG1113 | CG6805 | CG32817 | | | | |
| rdgA | CG9008 | | | | | |
| CG32280 | | | | | | |
| lncRNA:CR45959 | | | | | | |
| bark | | | | | | |
| CG13284 | | | | | | |
| CG9095 | | | | | | |
| CG2781 | | | | | | |
| ko | | | | | | |
| GILT2 | | | | | | |
| CG11134 | | | | | | |
| Ac13E | | | | | | |
| Pgant2 | | | | | | |

### Female DEGs by Zone including overall

| PSC | PSC and CC | CC | MZ | MZ and CCs | PSC and MZ | PSC and MZ and CC | PSC and MZ and overall | MZ and overall | overall | MZ and CC and overall | CC and overall | PSC and overall | PSC and CC and overall | PSC and CC and MZ and overall |
|---|---|---|---|---|---|---|---|---|---|---|---|---|---|---|
| CG12112 | CG3939 | CG17778 | CG2444 | CG11590 | | | CG15784 | CG2444 | CG4301 | Arpc3B | CG32816 | Cpr5C | | Karl |
| Idgf4 | | Gasp | Gclc | | | | CR43211 | Gclc | NimC1 | CG14434 | | | | MCTS1 |
| CG15365 | | CG30148 | CG34331 | | | | | CG34331 | sano | CG43133 | | | | lncRNA:CR40469 |
| CG14855 | | CG10527 | Atf3 | | | | | Atf3 | dpr14 | CG4593 | | | | |
| CG13369 | | alpha-Est4 | CG17982 | | | | | CG17982 | CG34054 | Sxl | | | | |
| form3 | | CG14629 | CG7322 | | | | | CG7322 | mthl4 | | | | | |
| CCKLR-17D3 | | CG14270 | Chrac-16 | | | | | Chrac-16 | Tsp42Ed | | | | | |
| snRNA:U2:14B | | lncRNA:CR45530 | hpRNA:CR32207 | | | | | hpRNA:CR32207 | prage | | | | | |
| CG9220 | | CrebB | rdgB | | | | | rdgB | Rip11 | | | | | |
| Hers | | Galphai | lncRNA:CR42868 | | | | | lncRNA:CR42868 | CG9981 | | | | | |
| CG31183 | | Tep1 | lncRNA:CR30009 | | | | | lncRNA:CR30009 | Sox15 | | | | | |
| septin 4 | | CG7453 | CG12056 | | | | | CG12056 | CG1840 | | | | | |
| CG17321 | | | CG17109 | | | | | CG17109 | lawc | | | | | |
| Dl | | | Fas3 | | | | | Fas3 | png | | | | | |
| Vps2 | | | Aven | | | | | Aven | RhoGAP18B | | | | | |
| comm2 | | | lncRNA:CR43264 | | | | | lncRNA:CR43264 | | | | | | |
| NimB5 | | | Dab | | | | | Dab | | | | | | |
| lncRNA:CR33218 | | | lncRNA:CR45517 | | | | | lncRNA:CR45517 | | | | | | |
| CG10508 | | | Wsck | | | | | Wsck | | | | | | |
| TkR99D | | | CG31689 | | | | | CG31689 | | | | | | |
| CG14516 | | | CR43217 | | | | | CR43217 | | | | | | |
| CG8369 | | | His2B:CG17949 | | | | | His2B:CG17949 | | | | | | |
| lncRNA:CR44933 | | | RNaseP:RNA | | | | | RNaseP:RNA | | | | | | |
| comm | | | | | | | | | | | | | | |
| hlk | | | | | | | | | | | | | | |
| asRNA:CR45912 | | | | | | | | | | | | | | |
| Ace | | | | | | | | | | | | | | |
| Dh31-R | | | | | | | | | | | | | | |
| CG17121 | | | | | | | | | | | | | | |
| mthl7 | | | | | | | | | | | | | | |
| CG34002 | | | | | | | | | | | | | | |
| CG6287 | | | | | | | | | | | | | | |
| ImpL2 | | | | | | | | | | | | | | |
| CG43759 | | | | | | | | | | | | | | |

### Male DEGs by Zone including overall

| PSC | PSC and CC | CC | MZ | MZ and CCs | PSC and MZ | PSC and MZ and CC | PSC and MZ and overall | MZ and overall | overall | MZ and CC and overall | CC and overall | PSC and overall | PSC and CC and overall | PSC and CC and MZ and overall |
|---|---|---|---|---|---|---|---|---|---|---|---|---|---|---|
| CG12112 | CG3939 | CG17778 | CG2444 | CG11590 | | | CG15784 | CG2444 | CG4301 | Arpc3B | CG32816 | Cpr5C | | Karl |
| Idgf4 | | Gasp | Gclc | | | | CR43211 | Gclc | NimC1 | CG14434 | | | | MCTS1 |
| CG15365 | | CG30148 | CG34331 | | | | | CG34331 | sano | CG43133 | | | | lncRNA:CR40469 |
| CG14855 | | CG10527 | Atf3 | | | | | Atf3 | dpr14 | CG4593 | | | | |
| CG13369 | | alpha-Est4 | CG17982 | | | | | CG17982 | CG34054 | Sxl | | | | |
| form3 | | CG14629 | CG7322 | | | | | CG7322 | mthl4 | | | | | |
| CCKLR-17D3 | | CG14270 | Chrac-16 | | | | | Chrac-16 | Tsp42Ed | | | | | |
| snRNA:U2:14B | | lncRNA:CR45530 | hpRNA:CR32207 | | | | | hpRNA:CR32207 | prage | | | | | |
| CG9220 | | CrebB | rdgB | | | | | rdgB | Rip11 | | | | | |
| Hers | | Galphai | lncRNA:CR42868 | | | | | lncRNA:CR42868 | CG9981 | | | | | |
| CG31183 | | Tep1 | lncRNA:CR30009 | | | | | lncRNA:CR30009 | Sox15 | | | | | |
| septin 4 | | CG7453 | CG12056 | | | | | CG12056 | CG1840 | | | | | |
| CG17321 | | | CG17109 | | | | | CG17109 | lawc | | | | | |
| Dl | | | Fas3 | | | | | Fas3 | png | | | | | |
| Vps2 | | | Aven | | | | | Aven | RhoGAP18B | | | | | |
| comm2 | | | lncRNA:CR43264 | | | | | lncRNA:CR43264 | | | | | | |
| NimB5 | | | Dab | | | | | Dab | | | | | | |
| lncRNA:CR33218 | | | lncRNA:CR45517 | | | | | lncRNA:CR45517 | | | | | | |
| CG10508 | | | Wsck | | | | | Wsck | | | | | | |
| TkR99D | | | CG31689 | | | | | CG31689 | | | | | | |
| CG14516 | | | CR43217 | | | | | CR43217 | | | | | | |
| CG8369 | | | His2B:CG17949 | | | | | His2B:CG17949 | | | | | | |
| lncRNA:CR44933 | | | RNaseP:RNA | | | | | RNaseP:RNA | | | | | | |
| comm | | | | | | | | | | | | | | |
| hlk | | | | | | | | | | | | | | |
| asRNA:CR45912 | | | | | | | | | | | | | | |
| Ace | | | | | | | | | | | | | | |
| Dh31-R | | | | | | | | | | | | | | |
| CG17121 | | | | | | | | | | | | | | |
| mthl7 | | | | | | | | | | | | | | |
| CG34002 | | | | | | | | | | | | | | |
| CG6287 | | | | | | | | | | | | | | |
| ImpL2 | | | | | | | | | | | | | | |
| CG43759 | | | | | | | | | | | | | | |
